# Microbead-Based Heterodimer Assay for Amplification-Free Detection of Single-Stranded DNA

**DOI:** 10.64898/2026.10.28.762408

**Authors:** Sona Rani Roy, Nicolas Lobato-Dauzier, Jeremy Mateos, Yannick Tauran, Arnaud Brioude, Alexandre Baccouche, Teruo Fujii, Anthony J. Genot, Soo Hyeon Kim

**Affiliations:** LIMMS/CNRS-Institute of Industrial Science, The University of Tokyo, 4-6-1 Komaba, Meguro-ku, Tokyo 153-8505, Japan; Sorbonne Université, CNRS, Institut de Biologie Paris-Seine, Laboratoire Jean Perrin, 75005, Paris, France; I3S Laboratory, Côte d’Azur University, CNRS, UMR 7271, Sophia Antipolis, France; Université Lyon 1, CNRS, LMI, UMR 5615, Villeurbanne, France; Institut d’Électronique, de Microélectronique et de Nanotechnologie, UMR 8520, Lille, France; Institute of Industrial Science, The University of Tokyo, 4-6-1 Komaba, Meguro-ku, Tokyo 153-8505, Japan

**Keywords:** nucleic acid detection, amplification-free detection, DNA hybridization, probe-functionalized microbeads, fluorescence microscopy

## Abstract

Direct detection of rare nucleic-acid targets without molecular amplification requires specific recognition events to be made observable while remaining distinguishable from nonspecific interactions. Here, we introduce a microbead-based heterodimer assay (MBHA) that converts two-site recognition of single-stranded DNA into spatially resolved, two-color bead-pairing events for direct nucleic-acid detection. Two spectrally distinct microbead populations are functionalized with partially double-stranded DNA probes exposing target-binding domains complementary to separate regions of the target. Target-mediated bridging produces heterotypic bead pairs that are classified by fluorescence identity and an interparticle-distance criterion, while residual background is evaluated relative to blank measurements. Under the tested conditions, 166 fM was the lowest target concentration exceeding the blank-derived detection threshold. A sequence-scrambled synthetic non-target DNA remained below threshold across the tested concentration range, and detection at 166 fM was retained in the presence of 1 µM non-target DNA, corresponding to an approximately six-million-fold molar excess. Probe redesign further extended the assay to a second 60-nucleotide synthetic DNA target derived from the SARS-CoV-2 ORF1ab sequence. This proof-of-concept study establishes heterodimer counting as an amplification-free measurement strategy in which rare molecular-recognition events are made directly observable and individually classifiable without increasing target copy number.

---

Molecular amplification underpins many of the most sensitive nucleic-acid assays. Polymerase chain reaction (PCR) and CRISPR–Cas-based methods provide high analytical sensitivity and sequence specificity for disease diagnostics and pathogen monitoring.^1–6^ Their implementation, however, commonly relies on enzymes, controlled reaction steps, and dedicated instrumentation, which can complicate decentralized testing.^7,8^ Amplification-free detection offers a complementary measurement regime in which recognition events are not multiplied through copy-number gain. Its central challenge is therefore to make rare target-mediated events directly observable while reliably distinguishing them from nonspecific interactions.

Probe-functionalized microbeads are well suited to this challenge because molecular binding can be translated into particle-scale optical or electrical signals.^9,10^ Existing platforms include multilayer fluorescent labeling,^11^ dielectrophoretic impedance measurement of bead-bound PCR products,^12^ bead-assisted target capture followed by real-time PCR,^13^ direct fluorescence imaging,^14^ flow-cytometric detection of bead-associated targets or aggregates,^15–17^ and paper-based aggregation readouts.^18^ These approaches span both amplified and direct measurement regimes.^11–18^ Target-mediated aggregation provides a further particle-scale readout, in which nucleic acids bridge probe-functionalized beads.^16,18,19^ Sloane et al. used hybridization-induced aggregation for KRAS mutation discrimination after PCR,^16,19^ whereas Kalish et al. converted target-mediated microsphere aggregation into a paper-based wicking-distance response that required target concentrations in the nM–µM range.^18^ Sun et al. further showed that auxiliary DNA hybridization and magnetic facilitation can promote flow-cytometric immunoaggregation for protein detection.^17^ Together, these studies establish aggregation as a useful transduction mechanism, but their readouts generally infer target binding from aggregate extent or a bulk response rather than classify heterotypic bead contacts. Distinguishing rare target-mediated contacts from nonspecific bead associations therefore remains a central challenge for direct aggregation-based detection.

Here, we introduce a microbead-based heterodimer assay (MBHA), an amplification-free particle-level strategy for direct nucleic-acid detection that distinguishes target-mediated heterotypic bead pairing from nonspecific associations. MBHA combines two-site molecular recognition with fluorescence identity, centroid-to-centroid proximity, and a blank-derived event-frequency threshold. The assay architecture and image-analysis workflow are summarized in Figure 1. Two spectrally distinct bead populations are functionalized with partially double-stranded DNA probes designed to expose single-stranded domains complementary to separate regions of the target (Figure 1a, steps i and ii). Target hybridization bridges the two bead types to produce heterotypic pairs (Figure 1a, step iii). Rather than using total aggregation as the analytical signal, MBHA classifies candidate events by fluorescence identity and an interparticle-distance criterion, excludes same-color homodimers, and evaluates event frequency relative to blank measurements (Figure 1b). The analytical logic therefore combines two-site molecular recognition, heterotypic fluorescence pairing, geometric proximity, and a sample-level event frequency exceeding the corresponding blank-derived threshold. We separately examine bead density and mixing duration as operational parameters affecting productive encounters and nonspecific contacts. Under the tested conditions, 166 fM was the lowest target concentration exceeding the blank-derived threshold. Notably, a sequence-scrambled synthetic non-target DNA remained below threshold across the tested concentration range, and detection at 166 fM was retained in the presence of 1 µM of the same non-target DNA, corresponding to an approximately six-million-fold molar excess. Probe redesign further extended the architecture to a second 60-nucleotide synthetic DNA target derived from the SARS-CoV-2 ORF1ab sequence. Together, these results position MBHA as a proof-of-concept amplification-free strategy that makes rare target-mediated interactions directly observable through classifiable particle-scale events, without increasing target copy number.

**Figure 1.**
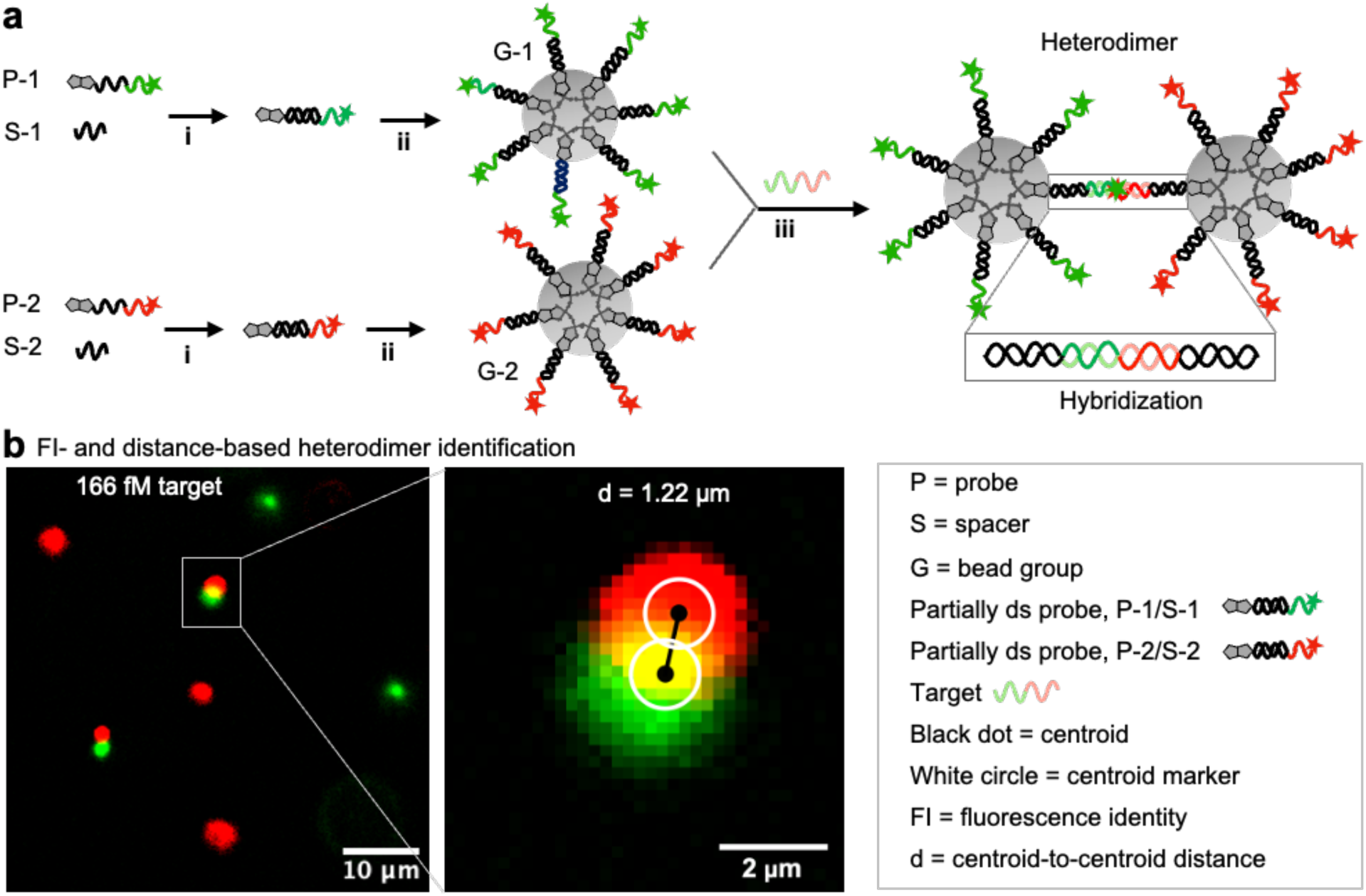
Working principle of MBHA and heterodimer identification. **(a)** Preparation of partially double-stranded, fluorescently labeled, dual-biotinylated probes P-1/S-1 and P-2/S-2 and their immobilization onto microbeads to generate G-1 and G-2, respectively, followed by target-mediated heterodimer formation. (i) P-1/S-1 and P-2/S-2 were annealed at 90 °C for 3 min followed by gradual cooling. (ii) Annealed probes were immobilized onto streptavidin-coated microbeads by gentle rotation overnight at room temperature. (iii) Heterodimer formation was promoted by mixing at 400 rpm for 30– 60 min followed by overnight incubation at room temperature. **(b)** Representative fluorescence microscopy image obtained with 166 fM target and enlarged example of heterodimer identification based on fluorescence identity (FI) and the centroid-to-centroid distance criterion. The representative green–red pair has a centroid-to-centroid separation of 5.53 pixels (1.22 µm). Black dots indicate the centroids, whereas white circles are graphical markers used to visualize the centroid positions. Scale bar in the fluorescence micrograph, 10 µm. Scale bar in the enlarged fluorescence micrograph, 2 µm.

## Experimental Section

### Oligonucleotides

All oligonucleotides were purchased from Integrated DNA Technologies (IDT) and Biomers.net (Table S1). Upon receipt, the oligonucleotides were dissolved in 1× TE buffer to a final concentration of 100 µM and stored at −20 °C.

Two probe sets, P-1/P-2 and P-3/P-4, were designed for two synthetic DNA targets. The first set, P-1 and P-2, consisted of 35-nt single-stranded oligonucleotides designed to hybridize to separate regions of the 40-nt model target T1′–2′. P-1 contained a 17-nt spacer region and an 18-nt target-binding region, whereas P-2 contained an 18-nt spacer region and a 17-nt target-binding region. Complementary spacer strands S-1 (17 nt) and S-2 (18 nt) were designed to form double-helical regions with P-1 and P-2, respectively. The target-binding regions of P-1 and P-2 were complementary to different regions of T1′–2′, allowing the target to bridge the two probes. P-1 was labeled with Atto488 at the 3′ end and P-2 with Atto647N at the 5′ end for fluorescence-based discrimination. Each probe carried two biotin groups at the terminus opposite its fluorophore for immobilization onto streptavidin-coated microbeads. A 70-nt scrambled sequence, Sc1′–2′, sharing approximately 50% sequence identity with T1′–2′, was used to evaluate sequence specificity.

The second probe set, P-3 and P-4, consisted of 60-nt oligonucleotides designed for a 60-nt synthetic DNA target, S3′–4′, derived from two regions of the SARS-CoV-2 ORF1ab sequence (GenBank accession no. PV239616.1). P-3 and P-4 each contained a 30-nt target-binding domain and a 30-nt spacer domain. In the P-3/P-1 construct, the 30-nt spacer domain of P-3 hybridized to a complementary 30-nt segment of P-1, leaving 5 nt of P-1 unpaired. Likewise, in the P-4/P-2 construct, the 30-nt spacer domain of P-4 hybridized to a complementary 30-nt segment of P-2, leaving 5 nt of P-2 unpaired. The exposed 30-nt target-binding domains of P-3 and P-4 were complementary to the two halves of S3′–4′. A 25-nt random oligonucleotide, RanN, was included as carrier DNA.^20^

### Microbead Functionalization

Streptavidin-coated polystyrene microbeads with a nominal diameter of 1 µm (Bangs Laboratories) were used. For each target, two probe-functionalized bead populations were prepared. For T1′–2′ detection, G-1 and G-2 were functionalized with partially double-stranded probes P-1/S-1 and P-2/S-2, respectively. For S3′–4′ detection, G-3 and G-4 were functionalized with partially double-stranded P-3/P-1 and P-4/P-2 complexes, respectively. In each second-target complex, a 30-nt spacer domain of P-3 or P-4 formed a 30-bp duplex with a complementary segment of the 35-nt biotinylated P-1 or P-2 strand, respectively. P-1/S-1 and P-2/S-2 were prepared by annealing P-1 and P-2 with S-1 and S-2, respectively. A nominal probe input of 3.85 × 10^5^ strands per bead was supplied in excess to promote high surface coverage.

According to the manufacturer’s certificate, the beads have a binding capacity of 0.63 µg biotin-FITC/mg of beads (1.56 × 10^12^ beads/g; 1.56 × 10^10^ beads/mL), corresponding to approximately 2.9 × 10^5^ biotin-binding sites per bead. Because the probes were dual biotinylated, this nominal strand input should not be interpreted as the actual number of immobilized probes per bead. Functionalization was performed in high-salt buffer (HSB; 2× TE, 1 M NaCl, and 0.2% Tween 20) to promote probe immobilization by reducing electrostatic repulsion between DNA and the bead surface. For G-1 and G-2, 6 µM of each probe (1.8 × 10^14^ strands) was annealed with 12 µM of the corresponding spacer oligonucleotide in 50 µL HSB. Annealing was performed at 90 °C for 3 min followed by gradual cooling to 30 °C at −3 °C per cycle. For G-3 and G-4, P-3 and P-4 were each used at 12 µM, whereas the corresponding P-1 and P-2 spacer strands were maintained at 6 µM to limit competition from their dual-biotin functionality during surface immobilization. During cooling, 30 µL of bead suspension was washed twice with 550 µL HSB by centrifugation at 5,000 × g for 4 min. The annealed probe-spacer complexes were then incubated with the washed beads in 600 µL HSB overnight at room temperature under gentle rotation. Unbound probes and spacers were removed by four washes with low-salt buffer (LSB; 1× TE, 50 mM NaCl, and 0.2% Tween 20). Functionalized beads were resuspended and stored in LSB supplemented with 2 mg/mL bovine serum albumin (BSA) to reduce nonspecific interactions and support bead dispersion.^21,22^

### Heterodimer Assay with Target DNA

Unless otherwise stated, heterodimer assays were performed in 50 µL LSB supplemented with 2 mg/mL BSA and 1 µM RanN. Spiking assays were performed in 25 µL. Standard 96-well plates with an approximately 40 mm2 well area were used for 50 µL assays, whereas half-area 96-well plates (PerkinElmer; approximately 20 mm2) were used for 25 µL assays. BSA served as a blocking agent to reduce nonspecific interactions,21,22 and RanN was included as carrier DNA to reduce sample loss due to adsorption.^20^

A master mix containing equal proportions of the two relevant bead populations (G-1/G-2 or G-3/G-4), BSA, and RanN was prepared. Either 48 µL of master mix for a 50 µL assay or 23 µL for a 25 µL assay was dispensed into each well. The plate was preincubated for 30 min at 300 rpm at room temperature to allow BSA and RanN to condition the well surface and reduce adsorption of target nucleic acids.^23^

To examine bead-density effects, three surface densities were tested: 1.25 × 10^3^, 2.5 × 10^3^, and 5 × 10^3^ microbeads/mm^2^. During bead preincubation, T1′–2′, Sc1′–2′, and S3′–4′ working solutions were prepared in Milli-Q water. All target concentrations and copy-number concentrations reported in this study refer to final in-well values after addition of a 2 µL target aliquot. For 50 µL assays, working solutions of 2.5 × 10^3^ to 2.5 × 10^8^ copies/µL were diluted 25-fold in the completed wells, giving 5 × 10^3^ to 5 × 10^8^ copies/well and final concentrations of 1 × 10^2^ to 1 × 10^7^ copies/µL. For 25 µL spiking assays, working solutions of 1.25 × 10^3^ to 1.25 × 10^8^ copies/µL were diluted 12.5-fold, giving 2.5 × 10^3^ to 2.5 × 10^8^ copies/well and the same final range of 1 × 10^2^ to 1 × 10^7^ copies/µL. Using 1 fM = 602.214 molecules/µL, these final concentrations correspond to 1.66 × 10^-1^, 1.66 × 10^0^, 1.66 × 10^1^, 1.66 × 10^2^, 1.66 × 10^3^, and 1.66 × 10^4^ fM. For the bead-density experiment, a 100-fold series was prepared analogously to give 1.66 × 10^-2^, 1.66 × 10^0^, 1.66 × 10^2^, and 1.66 × 10^4^ fM (10^1^, 10^3^, 10^5^, and 10^7^ copies/µL). Sc1′–2′ was tested from 1.66 × 10^-1^ to 1.66 × 10^4^ fM in 10-fold increments. For the interference experiment, Sc1′–2′ was present at a final concentration of 1 µM with each T1′–2′ concentration. An equal volume of water was added to blank assays. Samples were mixed at 400 rpm for 5, 30, or 60 min, depending on the experiment, and then incubated overnight at room temperature.

Blank assays were performed with six replicates; each tested target condition was evaluated in a single assay well.

### Fluorescence Microscopy

After overnight incubation, the plate was directly used for imaging. Imaging was performed using a Nikon Eclipse Ti2 inverted fluorescence microscope equipped with an LED illumination source. Excitation wavelengths of 488 and 635 nm were used for Atto488 and Atto647N, respectively, with illumination intensities of 5% at 488 nm and 15% at 635 nm. For experiments with the 60-nt DNA target, the Atto647N channel was imaged using 660 nm excitation at 15% illumination intensity. Images were acquired in multichannel scanning mode using a 20× objective and an exposure time of 100 ms.

Raw images were recorded in ND2 format and converted to TIFF files (3200 × 3200 pixels) using ImageJ. The TIFF files were used for heterodimer quantification in Mathematica. For presentation, images were cropped to 800 × 800 pixels (25% of the original width and height) for the Supporting Information and to 250 × 250 pixels (approximately 8% of the original width and height) for the main-text figures.

### Heterodimer Quantification of Microscopy Images

Heterodimer quantification was performed using Wolfram Mathematica with a custom image-analysis script (Mathematica Code S1). Fluorescence microscopy images were imported in TIFF format, and the green and red fluorescence images were processed independently. Fluorescent bead-like components were segmented using erosion, local-maximum detection, and connected-component analysis. For each segmented component, the centroid was determined from the coordinates of all pixels belonging to that component. For a component containing *N* pixels at coordinates (*xᵢ*, *yᵢ*), the centroid coordinates were calculated as

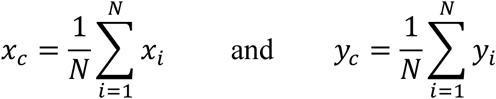

For each red-fluorescent component, the 30 nearest green-fluorescent components were identified. The centroid-to-centroid Euclidean distance between a green component at (*x*_1_, *y*_1_) and a red component at (*x*_2_, *y*_2_) was calculated as

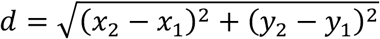

Candidate distance cutoffs of 6 and 8 pixels were evaluated because these values spanned the approximate centroid-to-centroid separation expected for associated bead pairs based on their apparent dimensions in the fluorescence images. For images acquired at 0.22 µm/pixel, these cutoffs corresponded to centroid-to-centroid distances of 1.32 and 1.76 µm, respectively. For one experimental dataset acquired at 0.33 µm/pixel, a 6-pixel cutoff corresponding to 1.98 µm was used. The cutoff was selected from the compact appearance of paired fluorescence objects in the processed images and was applied consistently within each experiment. The exact cutoff applied to each image analysis is reported in Tables S2–S6. A green-red pair was classified as a heterodimer association when its centroid-to-centroid distance was less than or equal to the specified cutoff. Only associations between differently labeled fluorescent components were included in the heterodimer signal. All qualifying green-red associations among the 30 nearest-neighbor candidates were counted.

The heterodimer abundance was normalized to the mean number of detected green- and red-fluorescent components and expressed as a percentage:

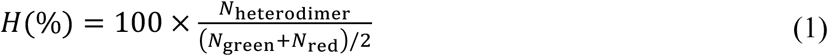

where *N*_heterodimer_ is the number of qualifying green–red associations, and *N*_green_ and *N*_red_ are the numbers of detected green- and red-fluorescent components, respectively. Normalization to the mean abundance of the two fluorescent populations accounts for differences in the numbers of green and red components detected in individual images. In aggregated structures, an individual bead could contribute to more than one counted heterotypic contact. Consequently, under conditions of extensive aggregation, the normalized heterodimer percentage may exceed 100%. In such cases, the value should be interpreted as a normalized heterotypic-contact metric rather than as the percentage of beads participating in unique one-to-one heterodimers.

To account for apparent heterodimer associations that may arise in blank samples from nonspecific bead association, probe interactions, or random spatial proximity, six blank measurements were used to establish the background response. The blank-derived reference threshold *T* was defined as the mean blank response plus three standard deviations:^24,25^

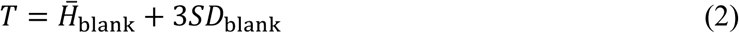

Where *H̄*_blank_ is the mean heterodimer percentage of the six blank measurements and *SD*_blank_ is the corresponding sample standard deviation. Thus, the reference threshold was positioned three standard deviations above the mean blank response.

The separation of each measured heterodimer signal from the blank distribution was additionally expressed as a Z-score:

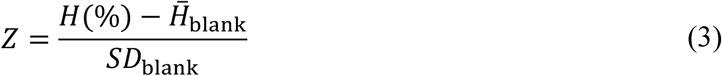

where *H* is the heterodimer percentage measured for the sample using eq 1. A Z-score of 3 indicates that the measured heterodimer percentage is three standard deviations above the mean blank response. Accordingly, *Z* ≥ 3 is mathematically equivalent to a heterodimer signal at or above the blank-derived threshold defined in eq 2 and was used as the criterion for positive detection.

## Results and Discussion

### Influence of Microbead Density on Background Signal and Target Detection

Microbead density can influence both nonspecific bead association and the probability of productive target-mediated encounters. We therefore examined three microbead densities, 1.25 × 10^3^, 2.5 × 10^3^, and 5 × 10^3^ microbeads/mm^2^, first in the absence of target to characterize background heterodimer formation and subsequently in the presence of T1′–2′ to determine whether bead density affected target detection. For both blank and target-containing assays, the microbead mixtures were agitated at 400 rpm for 60 min and then incubated overnight at room temperature. In target-containing assays, mixing was intended to promote bead-target encounters and hybridization, while the subsequent static incubation allowed further diffusion-mediated heterodimer formation^26–28^ and bead sedimentation for imaging.

In blank assays, the number of detected microbeads generally increased with bead density (Figure 2a, Figure S1a,c,e), although the increase was not proportional at the highest density, indicating nonuniform bead distribution across the well surface. More importantly, the mean nonspecific heterodimer percentage increased from 0.97% at 1.25 × 10^3^ microbeads/mm^2^ to 1.92% and 2.12% at 2.5 × 10^3^ and 5 × 10^3^ microbeads/mm^2^, respectively. Consequently, the blank-derived detection threshold increased from 2.36% to 3.45% and 3.60% across the same density range (Figure 2b, Table S2). These results indicate that higher microbead densities increase background heterotypic contacts, likely because reduced interparticle spacing increases opportunities for bead-bead contact and short-range nonspecific association.^29,30^

**Figure 2.**
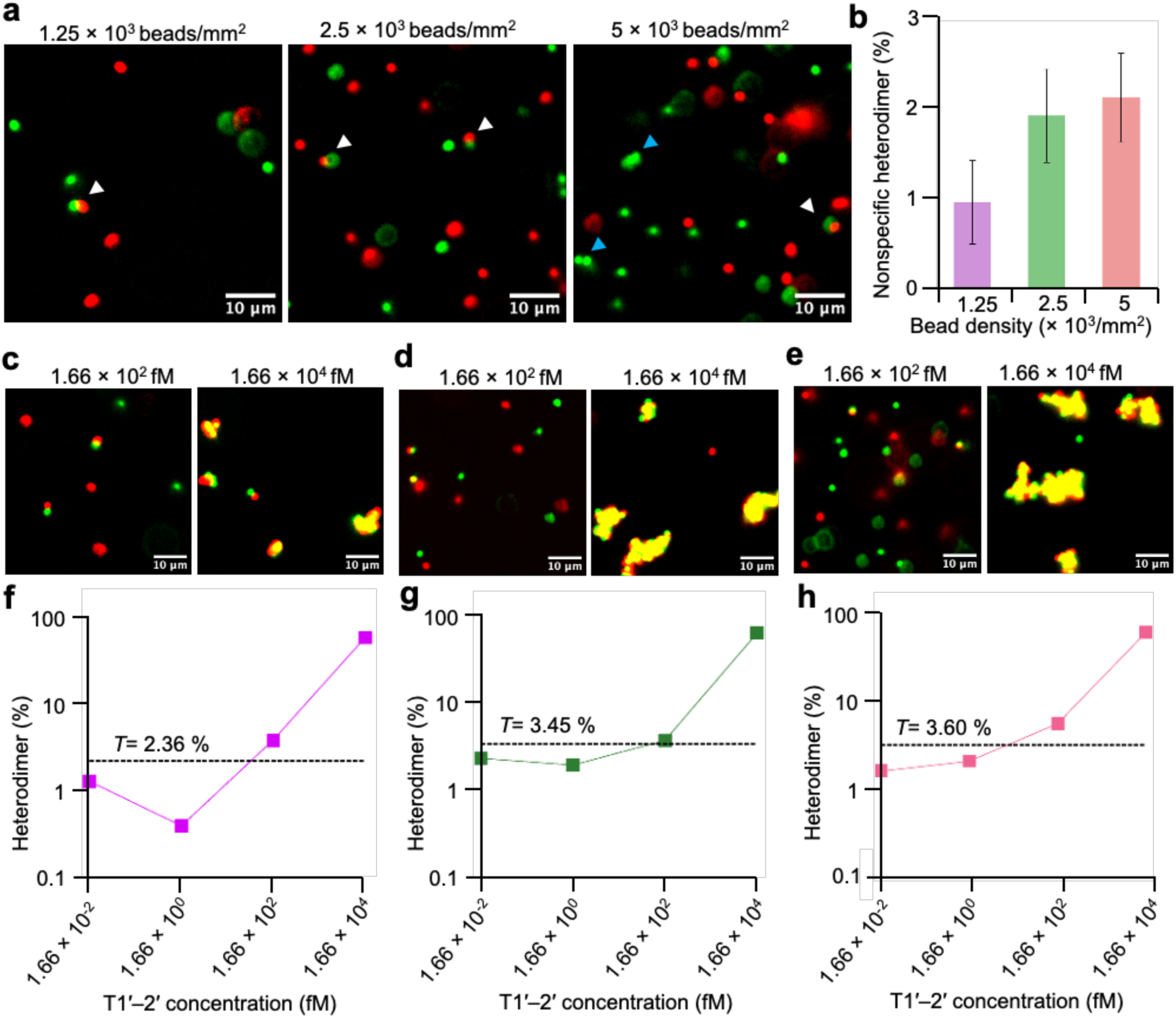
Influence of microbead density on background signal and target detection. **(a)** Representative fluorescence images of G-1/G-2 microbeads at densities of 1.25 × 10^3^, 2.5 × 10^3^, and 5 × 10^3^ microbeads/mm^2^ in blank assays. White arrowheads indicate heterodimer-like events, and blue arrowheads indicate homodimers. **(b)** Nonspecific heterodimer percentages measured at the three microbead densities. **(c–e)** Representative fluorescence images in the presence of T1′–2′ at microbead densities of **(c)** 1.25 × 10^3^, **(d)** 2.5 × 10^3^, and **(e)** 5 × 10^3^ microbeads/mm^2^. **(f–h)** Heterodimer percentages measured as a function of T1′–2′ concentration at microbead densities of **(f)** 1.25 × 10^3^**, (g)** 2.5 × 10^3^, and **(h)** 5 × 10^3^ microbeads/mm^2^. T1′–2′ concentrations were 1.66 × 10^-2^, 1.66 × 10^0^, 1.66 × 10^2^, and 1.66 × 10^4^ fM. *T* denotes the blank-derived detection threshold. Assays were performed in 50 µL with mixing at 400 rpm for 60 min, followed by overnight incubation. Blank assays comprised six replicates. Scale bars, 10 µm.

We next evaluated target-induced heterodimer formation using T1′–2′ concentrations of 1.66 × 10^-2^, 1.66 × 10^0^, 1.66 × 10^2^, and 1.66 × 10^4^ fM at the same three bead densities. Visible heterodimer clusters were observed at the highest target concentration across all densities, while lower concentrations did not produce visually distinguishable clusters (Figure 2c–e, Figure S1b,d,f). Larger aggregates were observed at higher bead densities, consistent with an increased probability of target-mediated bridging among neighboring beads.^31^ At the highest target concentration, extensive clustering also reduced the number of individually resolved beads (Table S2), indicating that absolute heterodimer percentages at this concentration should be interpreted primarily as a strong positive response rather than as a linear quantitative measure.

Quantitative analysis showed that 1.66 × 10^2^ fM T1′–2′ exceeded the corresponding blank-derived threshold at all three bead densities, with heterodimer percentages of 4.03%, 3.76%, and 6.41% for the low, intermediate, and high densities, respectively. At 1.66 × 10^4^ fM, the responses increased to 55.15%, 63.20%, and 68.60% for the low, intermediate, and high microbead densities, respectively (Figure 2f–h, Table S2). In contrast, the two lower target concentrations remained below threshold. The corresponding Z-scores showed the same classification because *Z* ≥ 3 is mathematically equivalent to exceeding a threshold defined as the blank mean plus three standard deviations (Table S2).

Overall, increasing microbead density enhanced both nonspecific background interactions and target-mediated heterodimer formation; however, it did not change the lowest concentration exceeding the detection threshold. T1′–2′ produced an above-threshold response at 166 fM across all three densities. Although higher bead densities increased the blank-derived threshold, increasing bead density provided no clear improvement in detection performance. The lowest tested density, 1.25 × 10^3^ microbeads/mm^2^, therefore provided the most favorable balance between minimizing background interactions and maintaining target detection and was selected for subsequent experiments.

### Effect of Mixing Duration on Heterodimer Formation

We next examined the effect of mixing duration on heterodimer formation at the selected microbead density of 1.25 × 10^3^ microbeads/mm^2^. T1′–2′ was tested from 1.66 × 10^-1^ to 1.66 × 10^4^ fM in 10-fold increments, and the bead suspension was mixed at 400 rpm for 5, 30, or 60 min before overnight static incubation. Blank measurements were obtained using the 60 min mixing condition.

Fluorescence images showed a relatively uniform microbead distribution, with visible heterodimer clusters appearing only at the highest target concentration across all three mixing durations (Figure 3a, Figure S2a–d). Across the assay series, mean detected bead counts ranged from 715 to 768 microbeads per field of view (Table S3). At the highest target concentration, the apparent bead count decreased, likely because compact aggregates were not fully resolved as individual beads.

**Figure 3.**
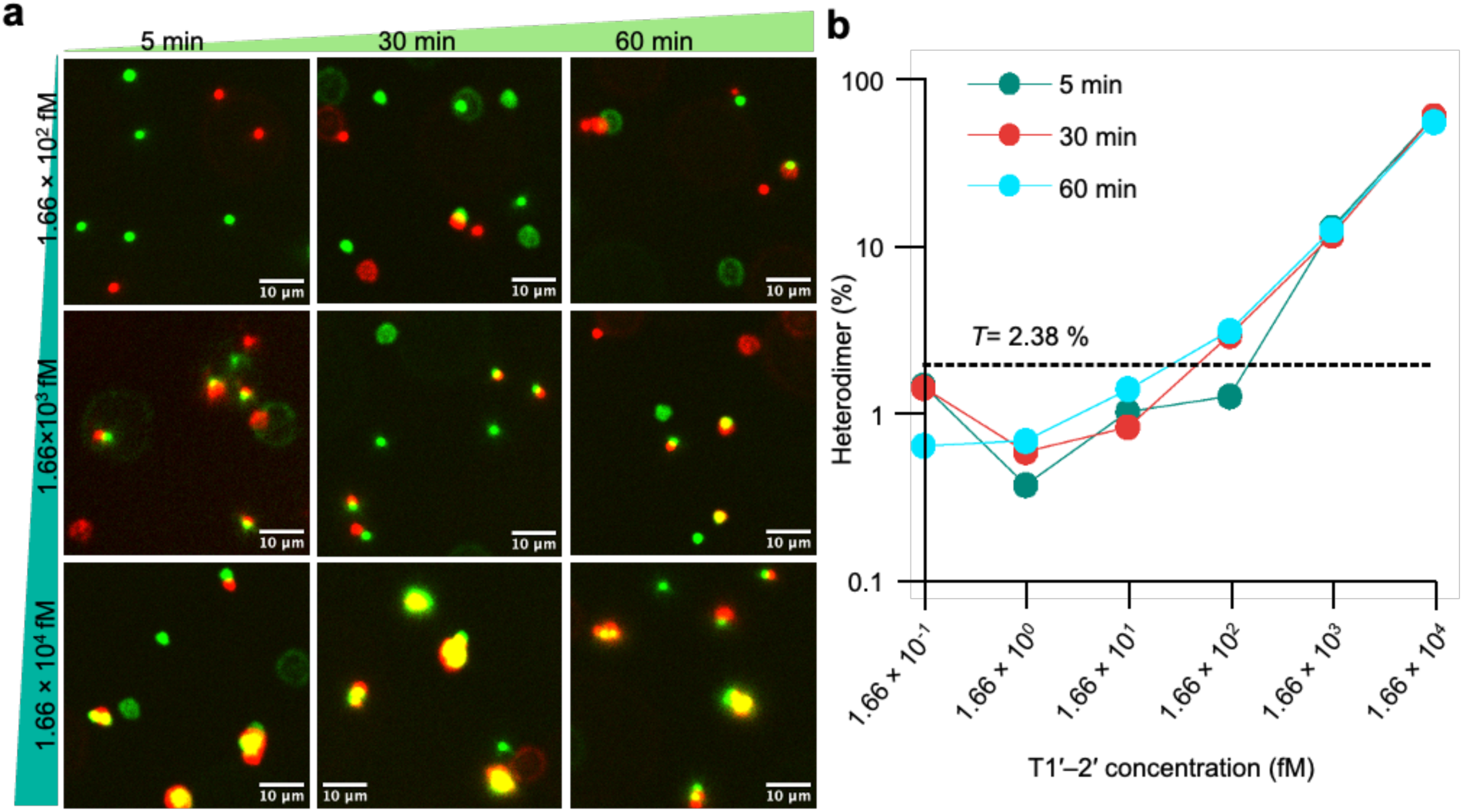
Effect of mixing duration on heterodimer formation. **(a)** Representative fluorescence images of G-1/G-2 microbeads and heterodimers at 1.66 × 10^2^, 1.66 × 10^3^, and 1.66 × 10^4^ fM T1′–2′ after 5, 30, or 60 min of mixing. Scale bars, 10 µm. **(b)** Heterodimer percentage as a function of T1′–2′ concentration for each mixing duration. Target concentrations ranged from 1.66 × 10^-1^ to 1.66 × 10^4^ fM in 10-fold increments. *T* denotes the blank-derived threshold. Bead density was 1.25 × 10^3^ microbeads/mm^2^ and assay volume was 50 µL. Blank assays comprised six replicates and were measured under the 60 min mixing condition.

The blank-derived detection threshold was 2.38%. After 5 min of mixing, heterodimer percentages of 14.41% and 61.93% were obtained at 1.66 × 10^3^ and 1.66 × 10^4^ fM, respectively, whereas lower concentrations were below threshold (Figure 3b). Extending the mixing duration to 30 or 60 min enabled detection at a 10-fold lower target concentration (Table S3). At 30 min, heterodimer percentages of 3.50%, 12.99%, and 62.40% were obtained at 1.66 × 10^2^, 1.66 × 10^3^, and 1.66 × 10^4^ fM, respectively. The corresponding values after 60 min were 3.72%, 13.92%, and 57.45%. Thus, both 30 min and 60 min mixing conditions produced above-threshold responses from 1.66 × 10^2^ fM (Figure 3b). The Z-score analysis gave the same detection classification (Table S3). Because blank measurements were performed only after 60 min of mixing, the same blank-derived threshold was used to evaluate all mixing durations. Therefore, this comparison does not show whether changing the mixing time itself affects background bead association.

These results indicate that 5 min of mixing may be insufficient, whereas 30 min of mixing improved detection by 10-fold. Increasing the mixing duration further to 60 min provided no additional improvement, suggesting that the benefit of active mixing largely plateaued by approximately 30 min. Although mechanical-force studies have shown that short DNA duplexes can be destabilized under applied force,^32,33^ the present experiments do not establish whether orbital mixing at 400 rpm affected bead-pair stability. Thus, 30 min of mixing was sufficient to promote the initial target-probe hybridization and bead-target interactions under the tested conditions.

Importantly, all samples subsequently underwent the same overnight static incubation. This step allowed continued diffusion-mediated target-probe hybridization and heterodimer formation while the microbeads gradually sedimented onto the imaging surface. Therefore, an adequate initial mixing period is important to promote initial interactions, whereas the subsequent static incubation provided additional time for bead pairing and sedimentation before imaging.

### Sequence Specificity and Detection in Excess Non-target DNA

The sequence specificity of MBHA was evaluated by comparing the heterodimer responses to T1′–2′ and the scrambled strand Sc1′–2′, which shares approximately 50% sequence identity with T1′–2′. Each sequence was tested separately from 1.66 × 10^-1^ to 1.66 × 10^4^ fM using G-1 and G-2 microbeads under the same heterodimer formation conditions.

In the presence of Sc1′–2′, fluorescence images remained similar to the blank samples across the full concentration range (Figure S3a–b), whereas T1′–2′ produced visible heterodimer clusters at the highest concentration (Figure 4a, Figure S3c). The blank-derived threshold was 1.60%, and all Sc1′–2′ measurements remained below this value. In contrast, T1′–2′ exceeded the threshold at 1.66 × 10^2^, 1.66 × 10^3^, and 1.66 × 10^4^ fM (Figure 4b). The corresponding Z-scores were above 3 at the same concentrations (Table S4), providing the equivalent standardized classification relative to the blank distribution.

**Figure 4.**
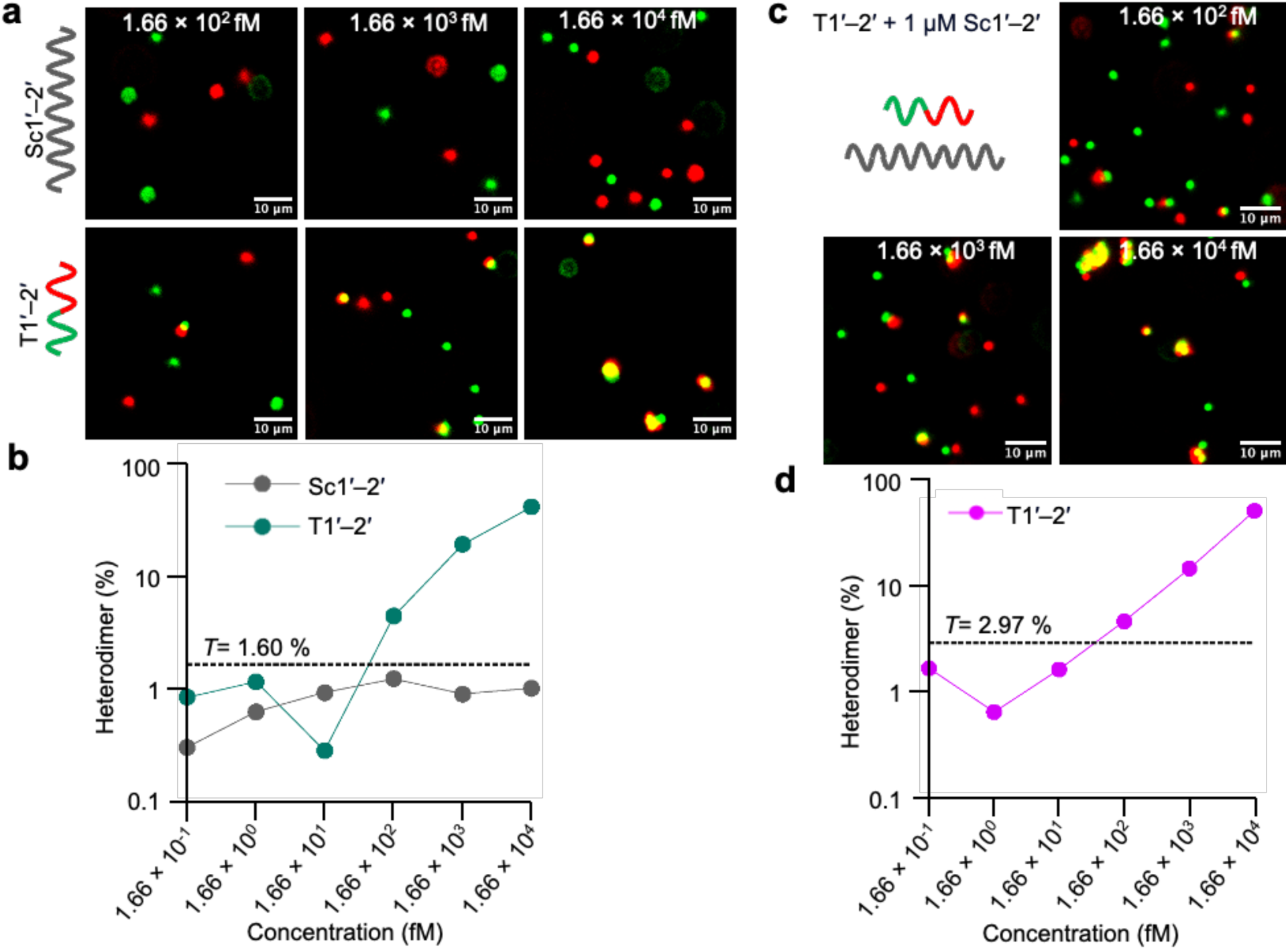
Sequence specificity and detection in excess non-target DNA. **(a)** Representative fluorescence images of G-1/G-2 microbeads with Sc1′–2′ or T1′–2′ at 1.66 × 10^2^, 1.66 × 10^3^, and 1.66 × 10^4^ fM. **(b)** Heterodimer percentages measured from 1.66 × 10^-1^ to 1.66 × 10^4^ fM. Assays were performed in 50 µL using a standard 96-well plate with mixing at 400 rpm for 60 min. **(c)** Spiking assay showing representative fluorescence images of T1′–2′ at 1.66 × 10^2^, 1.66 × 10^3^, and 1.66 × 10^4^ fM in the presence of 1 µM Sc1′–2′. **(d)** Heterodimer percentages measured from 1.66 × 10^-1^ to 1.66 × 10^4^ fM. Spiking assays were performed in 25 µL using a half-area 96-well plate with mixing at 400 rpm for 60 min. *T* denotes the blank-derived detection threshold. Blank assays comprised six replicates. Spiking blanks contained 1 µM Sc1′–2′. Scale bars, 10 µm.

To challenge target recognition in the presence of a large excess of non-target DNA, T1′–2′ was subsequently spiked into a background containing 1 µM Sc1′–2′. Under the spiking-assay conditions, visible heterodimer clusters were consistently observed at the highest target concentration (Figure 4c, Figure S4b). The blank-derived threshold was 2.97%. T1′–2′ remained above the blank-derived threshold at 1.66 × 10^2^, 1.66 × 10^3^, and 1.66 × 10^4^ fM, with heterodimer percentages of 4.73%, 14.80%, and 51.28%, respectively (Figure 4d, Table S5). The corresponding Z-scores also exceeded 3 at these concentrations, yielding the same classification relative to the blank distribution (Table S5). At 166 fM T1′–2′, the 1 µM Sc1′–2′ background corresponds to an approximately six-million-fold molar excess of the non-target strand. These results indicate sequence-selective heterodimer formation within the tested concentration range. Because the spiking assay used a different reaction volume and well format from the unspiked specificity experiment, the two datasets are best interpreted qualitatively rather than as a direct comparison of signal magnitude. The observed selectivity is consistent with preferential hybridization of complementary nucleic acid sequences under the assay conditions.^34,35^

### Detection of a Second SARS-CoV-2-Derived Synthetic DNA Target

To examine whether the MBHA architecture could be adapted to a second target sequence, we redesigned the probes for S3′–4′, a 60-nt synthetic DNA target derived from two regions of the SARS-CoV-2 ORF1ab sequence. Partially double-stranded P-3/P-1 and P-4/P-2 probes were immobilized onto G-3 and G-4 microbeads, respectively, and S3′–4′ was tested from 1.66 × 10^-1^ to 1.66 × 10^4^ fM using the selected low bead density and a 30 min mixing period (Figure 5a).

**Figure 5.**
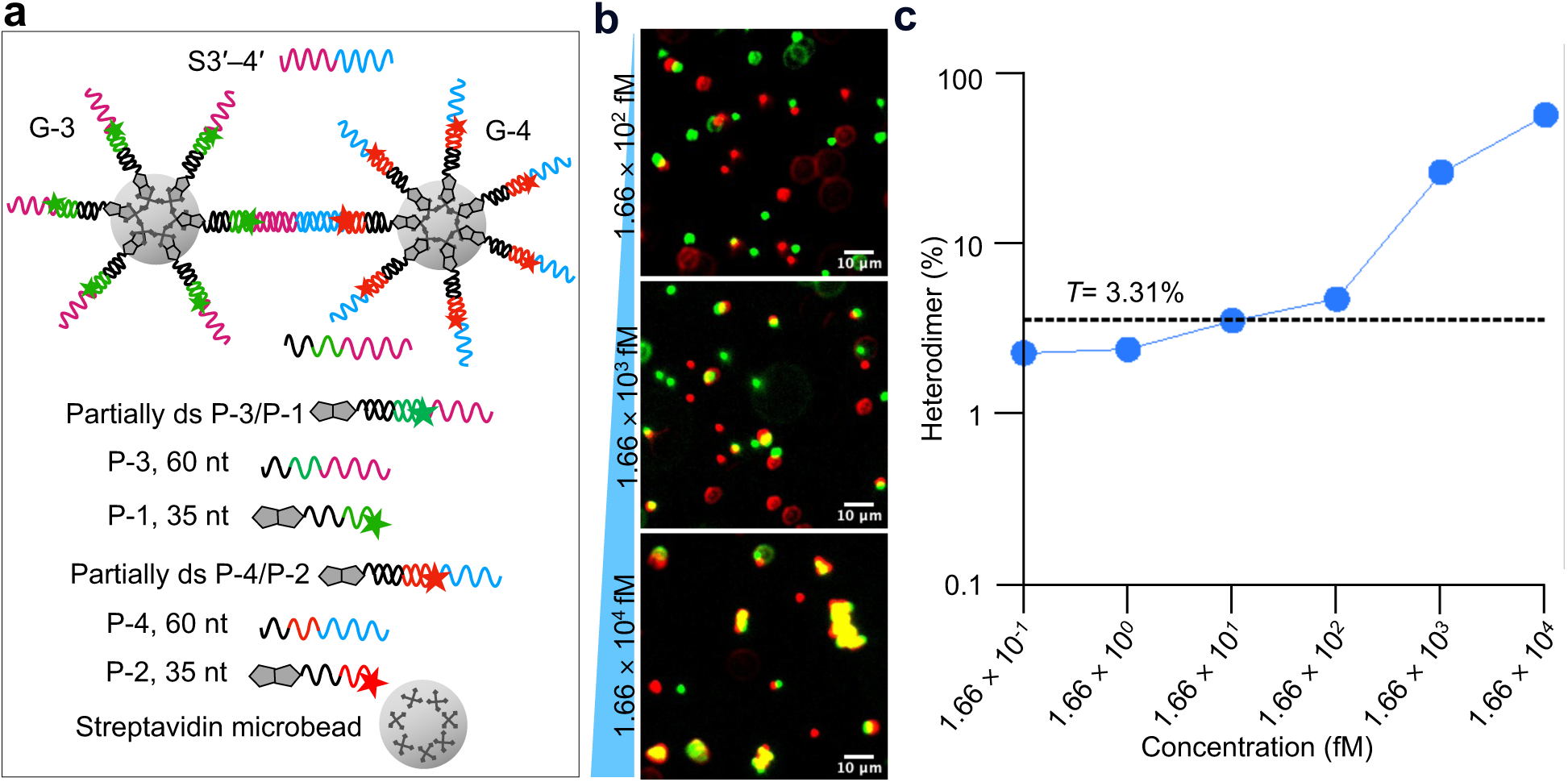
Detection of the 60-nt SARS-CoV-2 ORF1ab-derived synthetic DNA target S3′–4′. **(a)** Schematic representation of S3′–4′-mediated heterodimer formation using partially double-stranded P-3/P-1 and P-4/P-2 probe architectures on G-3 and G-4 microbeads, respectively. P-1 and P-2 are 35 nt; 30 nt of each strand participates in the corresponding duplex spacer. **(b)** Representative fluorescence images at 1.66 × 10^2^, 1.66 × 10^3^, and 1.66 × 10^4^ fM S3′–4′. **(c)** Heterodimer percentages were measured as a function of S3′–4′ concentration from 1.66 × 10^-1^ to 1.66 × 10^4^ fM. Mixing was performed at 400 rpm for 30 min. *T* denotes the blank-derived threshold. Blank assays comprised six replicates. Scale bars, 10 µm.

The S3′–4′ experiments showed a higher number of microbeads per field of view (Figure S5) than the preceding T1′–2′ experiments, with averages of 2,486 beads in blanks and 2,888 in target-containing samples (Table S6). Visible heterodimer clusters appeared only at the highest target concentration (Figure 5b, Figure S5b). This difference may reflect nonuniform bead distribution within the well or variation in the imaged field of view. The higher bead coverage was associated with a higher blank-derived threshold of 3.31% (Table S6) and should therefore be considered when comparing absolute heterodimer percentages across experiments.

Nevertheless, 166 fM remained the lowest tested concentration exceeding the blank-derived threshold, as in the T1′–2′ experiments.

Heterodimer percentages of 4.42%, 24.34%, and 52.61% were obtained at 1.66 × 10^2^, 1.66 × 10^3^, and 1.66 × 10^4^ fM S3′–4′, respectively, all exceeding the 3.31% threshold (Figure 5c). The corresponding Z-scores were also above 3 (Table S6). These results indicate that the partially double-stranded probe architecture can be redesigned for a second synthetic target while retaining an above-threshold response at 166 fM under the tested conditions. The observation also suggests that increasing probe length from the P-1/P-2 format to the P-3/P-4 format did not prevent target-mediated heterodimer formation.^26,36^

### Analytical Performance of MBHA

A key feature of MBHA is the direct relationship between its molecular recognition and signal-generation mechanism. Although nucleic-acid hybridization in biological environments can be influenced by molecular crowding, secondary structure, target accessibility, and interactions with other macromolecules,^37^ MBHA reduces target recognition to a defined two-site hybridization event in a relatively simple assay system using a defined laboratory buffer containing 50 mM NaCl. Simultaneous binding of a target to two complementary probe domains converts molecular recognition directly into a heterotypic bead-pairing event, consistent with the broader use of DNA hybridization to program specific interactions between colloidal particles.^31^ Importantly, this process requires no polymerase, nuclease, CRISPR-associated enzyme, or enzymatic reporter. The analytical signal arises directly from target hybridization and bead-pair classification, avoiding amplification- or enzyme-dependent signal generation.^38^

The partially double-stranded probe architecture provides an additional design feature for target recognition in MBHA. Each bead-bound probe contains a duplex spacer region and an exposed single-stranded target-binding domain. Related particle-based systems have used DNA constructs containing a double-stranded spacer with a distal single-stranded overhang to present accessible recognition sequences on streptavidin-coated particles.^39^ In addition, double-stranded linkers have been shown to improve the strength and kinetics of surface DNA hybridization at high probe densities, where electrostatic repulsion and steric hindrance can otherwise restrict target binding.^40^ In MBHA, the duplex spacer may therefore help position the single-stranded recognition domain away from the microbead surface and maintain its accessibility for hybridization. This arrangement is particularly relevant because a target must simultaneously engage complementary probe domains on two different bead populations to generate a heterotypic bead contact. Thus, the partially double-stranded probe design provides an exposed recognition configuration suitable for target-mediated bead bridging.

Across the T1′–2′ experiments, 166 fM was the lowest tested target concentration that produced a threshold-positive response under the present experimental conditions. Other highly sensitive nucleic-acid assays can achieve lower detection levels, but often rely on additional processing steps or specialized readout components.^38,41–44^ For example, SMOLT achieves sub-femtomolar detection using long DNA probes, functionalized magnetic beads and capillaries, controlled flow, washing, bead-displacement measurements, and a disruption step.^41^ Among amplification-free bead-based methods, reported detection limits include approximately 1.4 pM for magnetic modulation,^10^ 5 pM for magnetic-bead capture with fluorescent-probe imaging,^14^ and 200 pM for flow-cytometric fluorescence recovery.^15^ Under the present conditions, MBHA produced a threshold-positive response at 166 fM, a lower concentration than these reported values, while retaining a direct bead-based readout. Because the assay formats and detection principles differ, these values are best considered contextual rather than direct performance comparisons.

The spiking experiment further supports the analytical selectivity of MBHA under substantial sequence interference. Detection of 166 fM T1′–2′ in the presence of 1 µM Sc1′–2′, corresponding to an approximately six-million-fold molar excess of non-target DNA, indicates that target-mediated heterotypic bead contacts remained distinguishable from the blank response despite the large excess of non-target sequence. Related interference-challenge studies have been reported for other amplification-free nucleic-acid assays. Esfandiari et al. demonstrated 10 fM sequence-specific DNA detection against a background designed to simulate noncomplementary genomic DNA in real samples,^45^ while Máčala et al. showed target quantification in the presence of excess highly homologous non-target sequence using an amplification-free hybridization-complex-transfer assay.^46^ These amplification-free methods achieve lower detection limits through specialized transduction schemes or additional target-processing steps. For example, the hybridization-complex-transfer assay incorporates sequential capture, target-complex release, transfer, and recapture to suppress nonspecific background.^46^ In contrast, MBHA converts target recognition directly into heterotypic bead contacts classified by fluorescence identity and spatial proximity, without target amplification or post-hybridization washing. Because 166 fM was not established as a formal limit of detection in the present study, further replicated LOD analysis and validation in complex biological matrices will be required to establish analytical sensitivity and matrix tolerance under practical conditions.

Taken together, MBHA combines a threshold-positive response at 166 fM, sequence-selective recognition, and a comparatively direct signal-generation mechanism. Rather than relying on enzymatic amplification or multistep target-transfer procedures, MBHA converts sequence-specific hybridization directly into optically identifiable heterotypic bead contacts. The partially double-stranded probe architecture also provides a programmable recognition format, as demonstrated by redesign of the probe sequences for a second synthetic DNA target.

## Conclusions

We developed an enzyme-free microbead-based heterodimer assay (MBHA) that converts sequence-specific target bridging directly into classifiable heterotypic bead contacts. The assay requires no nucleic-acid amplification, polymerase, nuclease, CRISPR-associated enzyme, or enzymatic reporter. The analytical signal arises directly from complementary hybridization between two probe-functionalized bead populations. Under the present experimental conditions, 166 fM was the lowest tested target concentration that exceeded the blank-derived threshold.

Importantly, the spiking experiment demonstrated that target-specific heterodimer formation remained distinguishable from background even under substantial interference from the tested non-target sequence, further supporting the analytical selectivity of the assay.

The results also show that assay performance depends on controlling nonspecific bead interactions and providing sufficient time for productive bead-target encounters. The partially double-stranded probe architecture enabled the same recognition principle to be redirected to a second synthetic DNA target through probe redesign, supporting the adaptability of the platform. Together, these findings establish heterotypic bead-contact classification as a simple particle-level transduction strategy that differs from conventional bulk aggregation measurements by combining fluorescence identity with spatial proximity to identify target-mediated bead associations.

The present study was performed using short synthetic DNA targets, and target-positive conditions were evaluated in single assay wells. Therefore, independent replication, formal limit-of-detection validation, and testing in biological and other complex sample matrices will be important next steps. The overnight incubation and reliance on fluorescence microscopy are additional practical limitations that may be reduced through optimization of reaction kinetics, microfluidic formats, and compact imaging platforms. Nevertheless, the combination of a threshold-positive response at 166 fM, substantial tolerance to the tested non-target sequence interference, absence of enzymatic amplification, and straightforward probe redesign provides a basis for further development of MBHA as a simple and adaptable nucleic-acid detection platform.

## Supporting Information

Oligonucleotide sequences, additional fluorescence microscopy images, heterodimer quantification tables, and Mathematica image-analysis code (PDF).

## Data Availability

The data supporting the findings of this study are available in the article and its Supporting Information. The Mathematica image-analysis code is provided as Mathematica Code S1.

## Author Information

### Corresponding Authors

*Sona Rani Roy* − LIMMS/CNRS-Institute of Industrial Science, The University of Tokyo, 4-6-1 Komaba, Meguro-ku, Tokyo 153-8505, Japan; ORCID: 0000-0002-9442-9566;

*Soo Hyeon Kim* – LIMMS/CNRS-Institute of Industrial Science, The University of Tokyo, 4-6-1 Komaba, Meguro-ku, Tokyo 153-8505, Japan; Institute of Industrial Science, The University of Tokyo, 4-6-1 Komaba, Meguro-ku, Tokyo 153-8505, Japan; ORCID: 0000-0001-9842-9523;

### Authors

*Nicolas Lobato-Dauzier* − Sorbonne Université, CNRS, Institut de Biologie Paris-Seine, Laboratoire Jean Perrin, 75005, Paris, France; ORCID: 0009-0007-3790-9217;

*Jeremy Mateos* − I3S Laboratory, Côte d’Azur University, CNRS, UMR 7271, Sophia Antipolis, France; ORCID: 0009-0002-4864-3876;

*Yannick Tauran* − Université Lyon 1, CNRS, LMI, UMR 5615, Villeurbanne, France; ORCID: 0000-0002-0706-7799;

*Arnaud Brioude* − Université Lyon 1, CNRS, LMI, UMR 5615, Villeurbanne, France; ORCID: 0000-0002-8036-3973;

*Alexandre Baccouche* − LIMMS/CNRS-Institute of Industrial Science, The University of Tokyo, 4-6-1 Komaba, Meguro-ku, Tokyo 153-8505, Japan; Institut d’Électronique, de Microélectronique et de Nanotechnologie, UMR 8520, Lille, France; ORCID: 0009-0000-5476-9585;

*Teruo Fujii* − Institute of Industrial Science, The University of Tokyo, 4-6-1 Komaba, Meguro-ku, Tokyo 153-8505, Japan; ORCID: 0000-0001-7529-6238; *Anthony J. Genot* − LIMMS/CNRS-Institute of Industrial Science, The University of Tokyo, 4-6-1 Komaba, Meguro-ku, Tokyo 153-8505, Japan; ORCID: 0000-0001-7535-7432.

### Author Notes

† Deceased April 2025.

### Author Contributions

Conceptualization: S.R.R., N.L.D., Y.T., A.J.G., S.H.K.; data curation: S.R.R.; funding acquisition: A.J.G., S.H.K.; investigation: S.R.R., N.L.D., J.M.; methodology: S.R.R., A.J.G.; project administration: S.R.R., A.J.G., S.H.K.; resources: Ar.B., T.F., A.J.G., S.H.K.; supervision: S.R.R., A.J.G., S.H.K.; validation: S.R.R.; visualization: S.R.R.; writing – original draft: S.R.R.; writing – review & editing: S.R.R., N.L.D., Al.B., Y.T., T.F., S.H.K.

## Acknowledgments

This work was supported by the ANR Flash COVID-19 Project (Grant Number: ANR-20-COVI-0041), the CNRS Prematuration Project at LIMMS, and JSPS KAKENHI (Grant Number: JP20K21204).

The authors acknowledge the use of ChatGPT (OpenAI) for language refinement, reference-formatting assistance, and assistance with code formatting and troubleshooting. All scientific content, experimental data, analyses, interpretations, and conclusions were independently developed, reviewed, and verified by the authors, who take full responsibility for the content of this manuscript.

