## Supporting Information for "Microbead-Based Heterodimer Assay for Amplification-Free Detection of Single-Stranded DNA"

[†] Deceased April 2025.

**Table of Contents**

Figure S1. Fluorescence images of microbeads at three densities S2
Figure S2. Fluorescence images of microbeads at three mixing durations S8
Figure S3. Fluorescence images from probe specificity assays S13
Figure S4. Fluorescence images from the spiking assay S17
Figure S5. Fluorescence images of microbeads with S3′–4′ S20
Table S1. Oligonucleotides used in this study S23
Table S2. Heterodimer counts from three different microbead densities S24
Table S3. Heterodimer counts after 5, 30, and 60 min of mixing S25
Table S4. Heterodimer counts from probe specificity assays S26
Table S5. Heterodimer counts from spiking assays S27
Table S6. Heterodimer counts from S3′–4′ S28
Mathematica Code S1. Image-based microbead and heterodimer quantification script S29

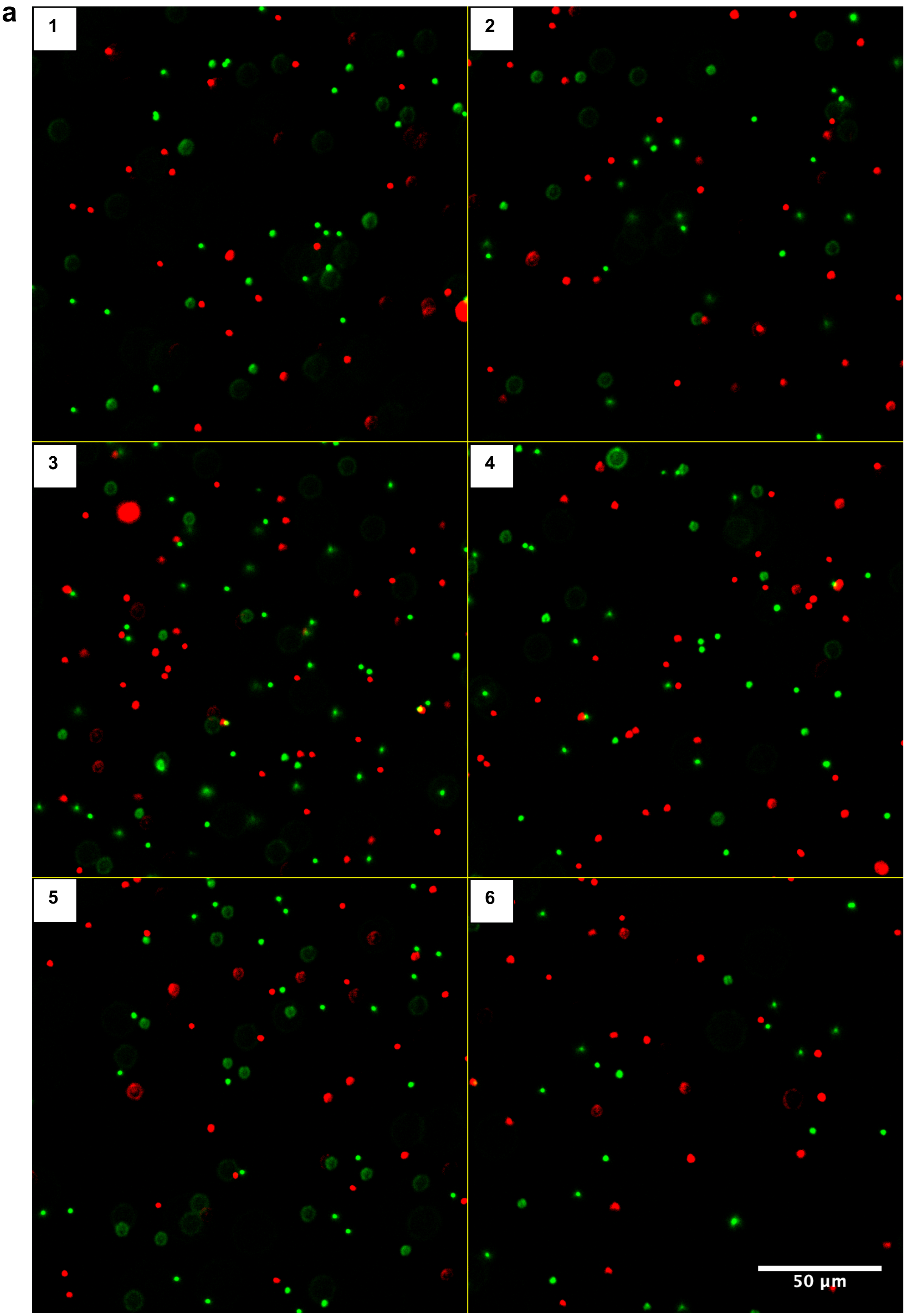
a
1
2
3
4
5
6
50 µm

b

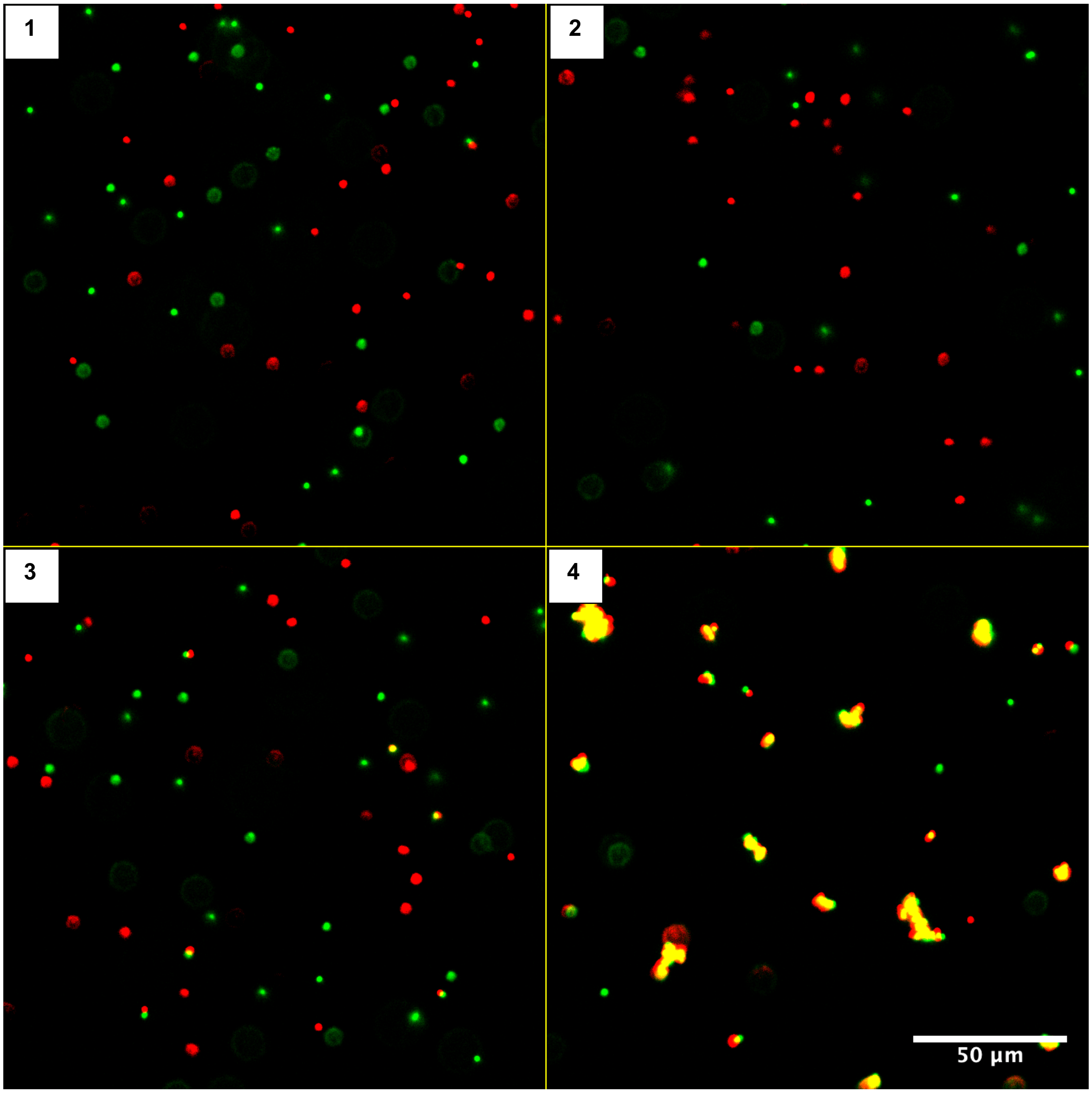

c

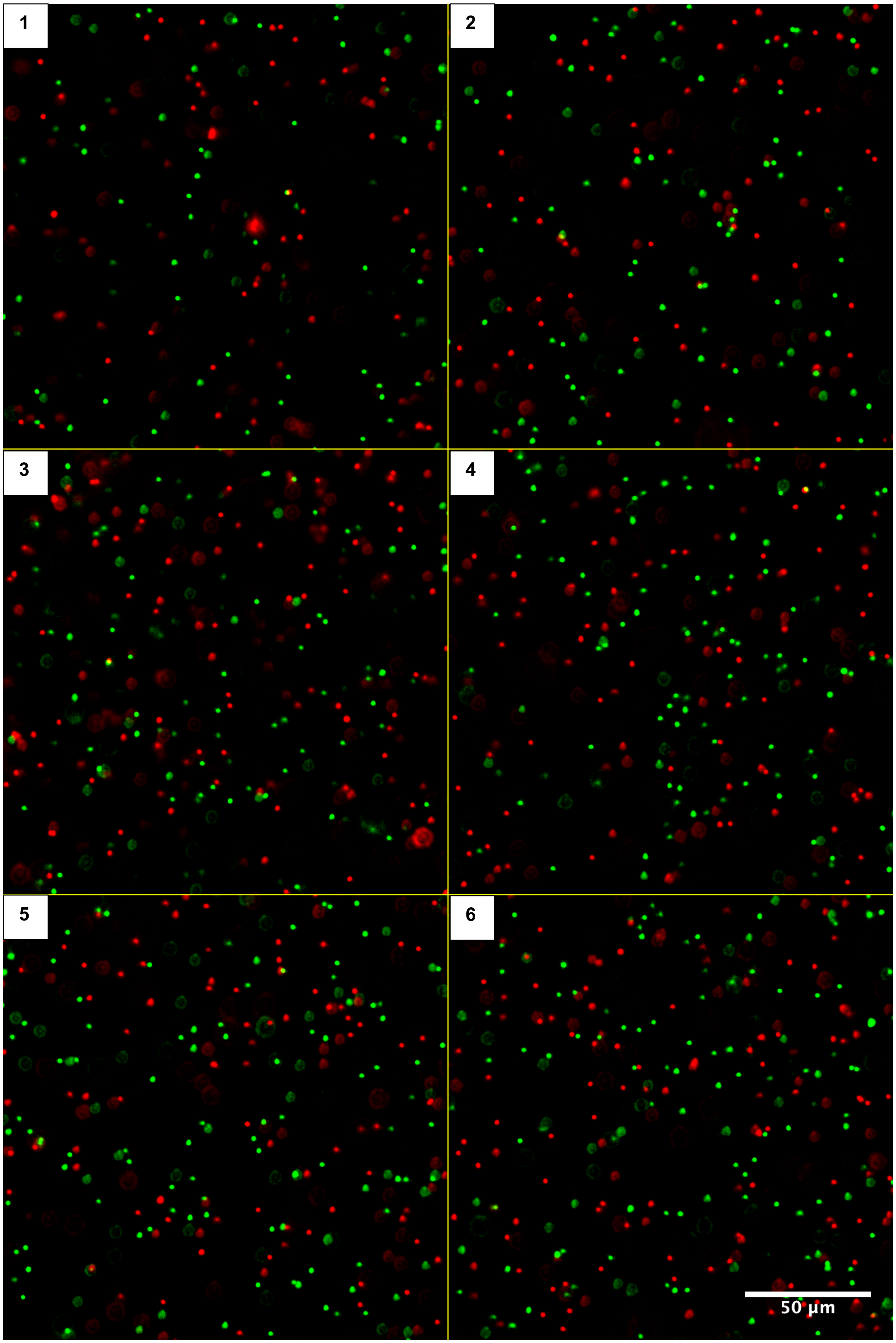

d

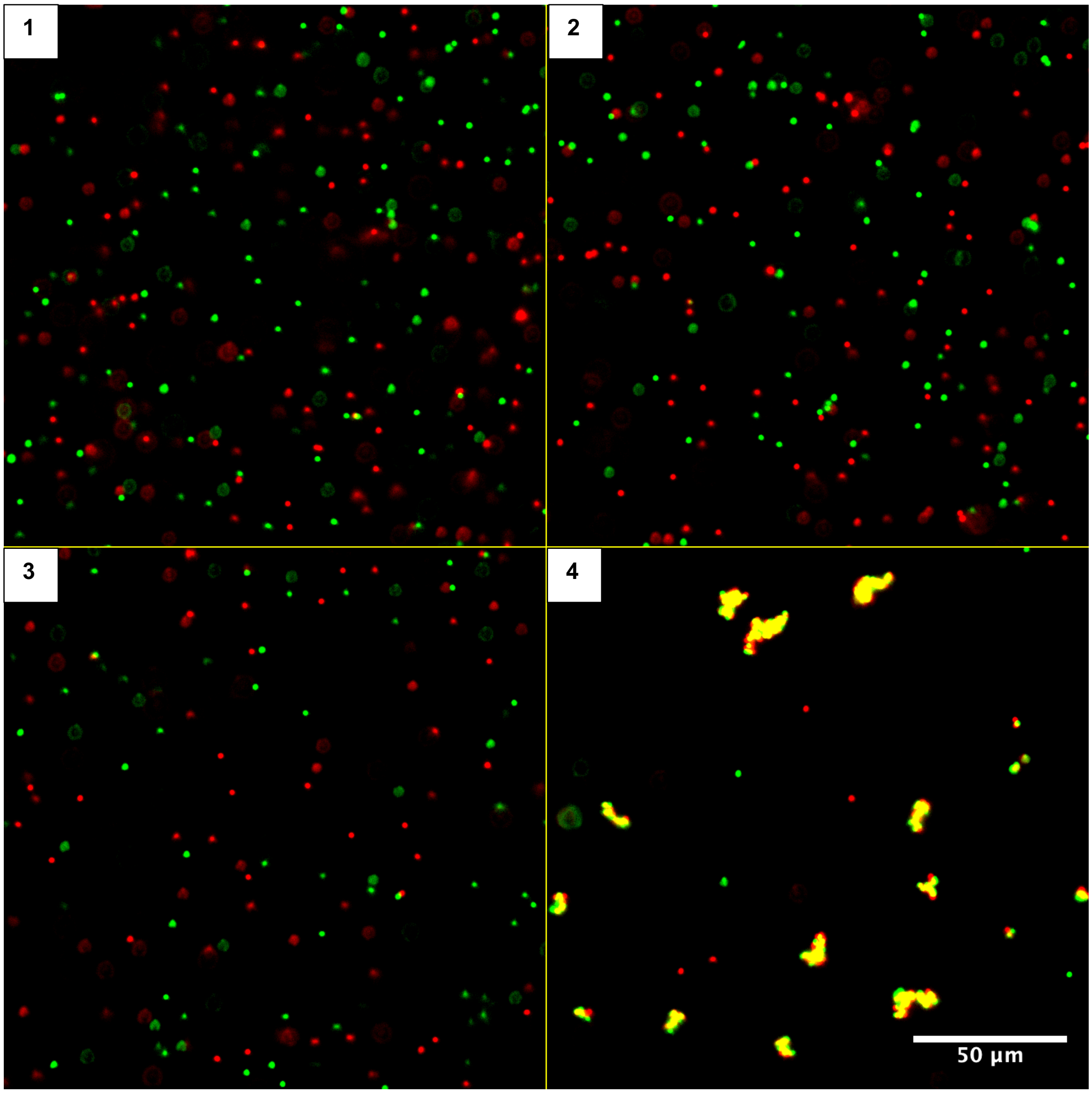

e

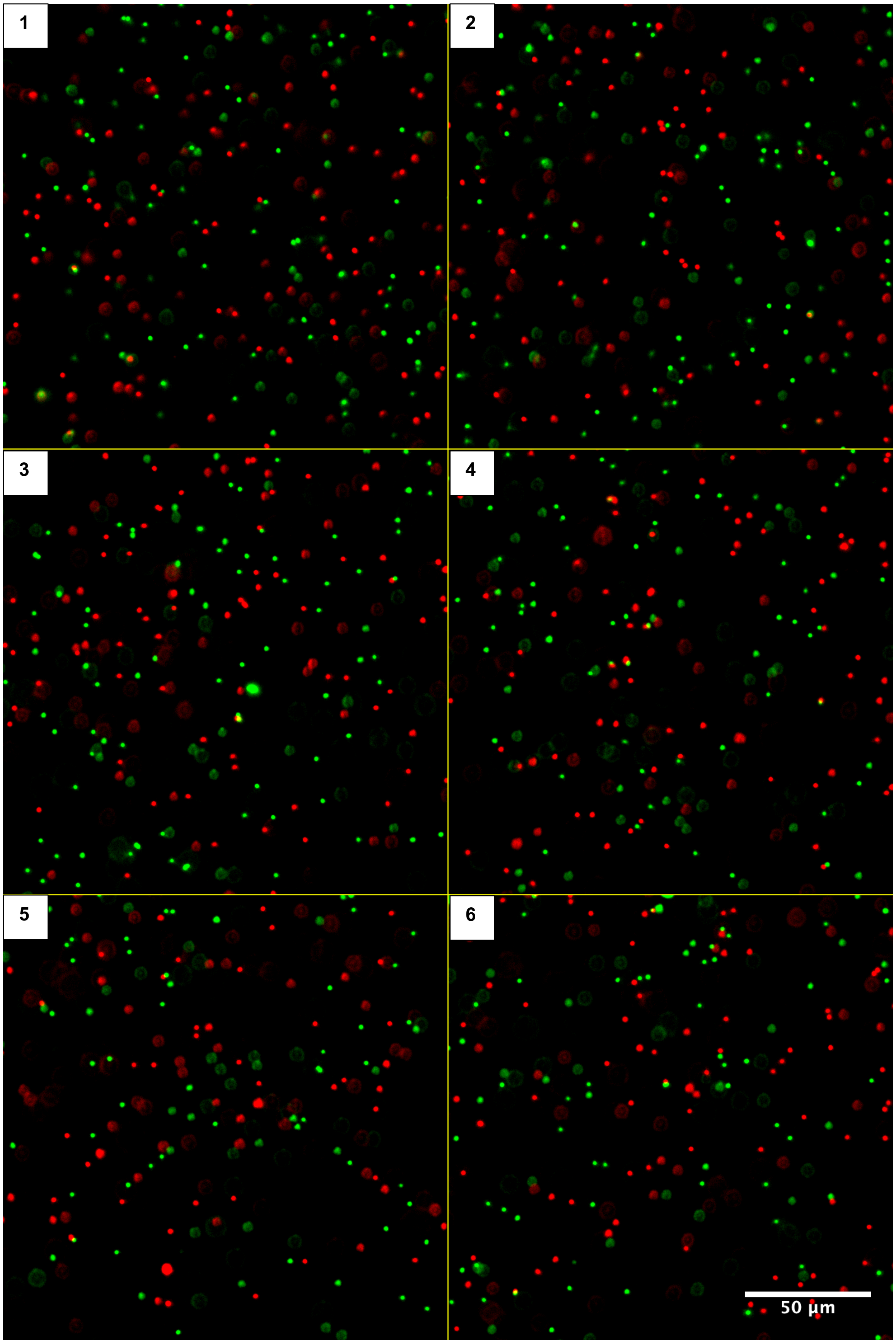

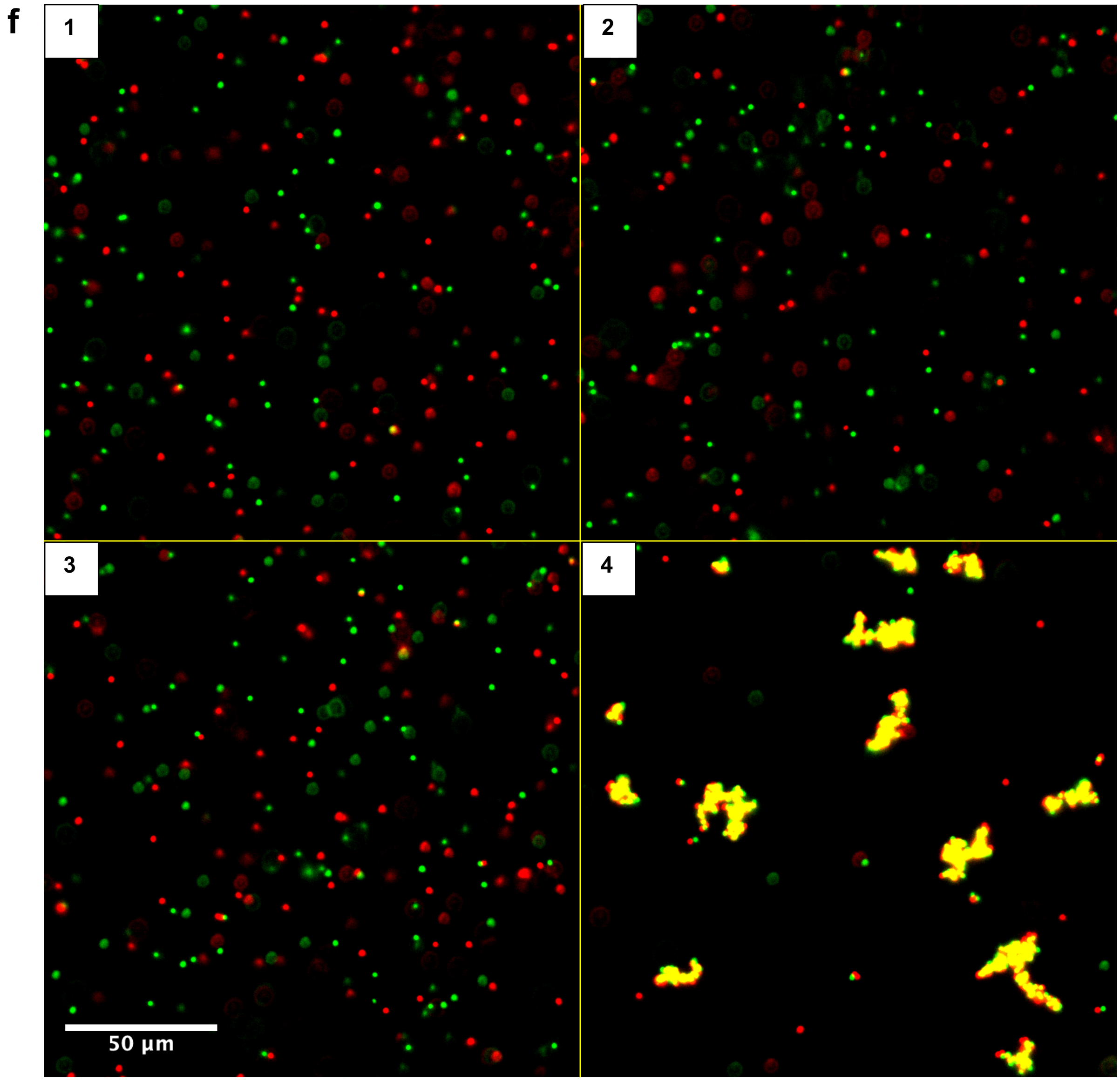


**Figure S1.** Fluorescence images of microbeads at three densities. **(a, b)** $1.25 \times 10^3$, **(c, d)** $2.5 \times 10^3$ and **(e, f)** $5 \times 10^3$ microbeads/mm$^2$, corresponding to approximately $5 \times 10^4$, $1 \times 10^5$, and $2 \times 10^5$ microbeads per well, respectively, based on an approximately 40 mm$^2$ well area. **(a, c, e)** Blank assays comprising six replicates (1–6) without target. **(b, d, f)** Assays with T1′–2′ at four concentrations: (1) $1.66 \times 10^{-2}$, (2) $1.66 \times 10^0$, (3) $1.66 \times 10^2$, and (4) $1.66 \times 10^4$ fM. Assays were performed in 50 µL with mixing at 400 rpm for 60 min. Images were cropped to 800 × 800 pixels (25% of the original width and height) for presentation. Scale bars, 50 µm.

a

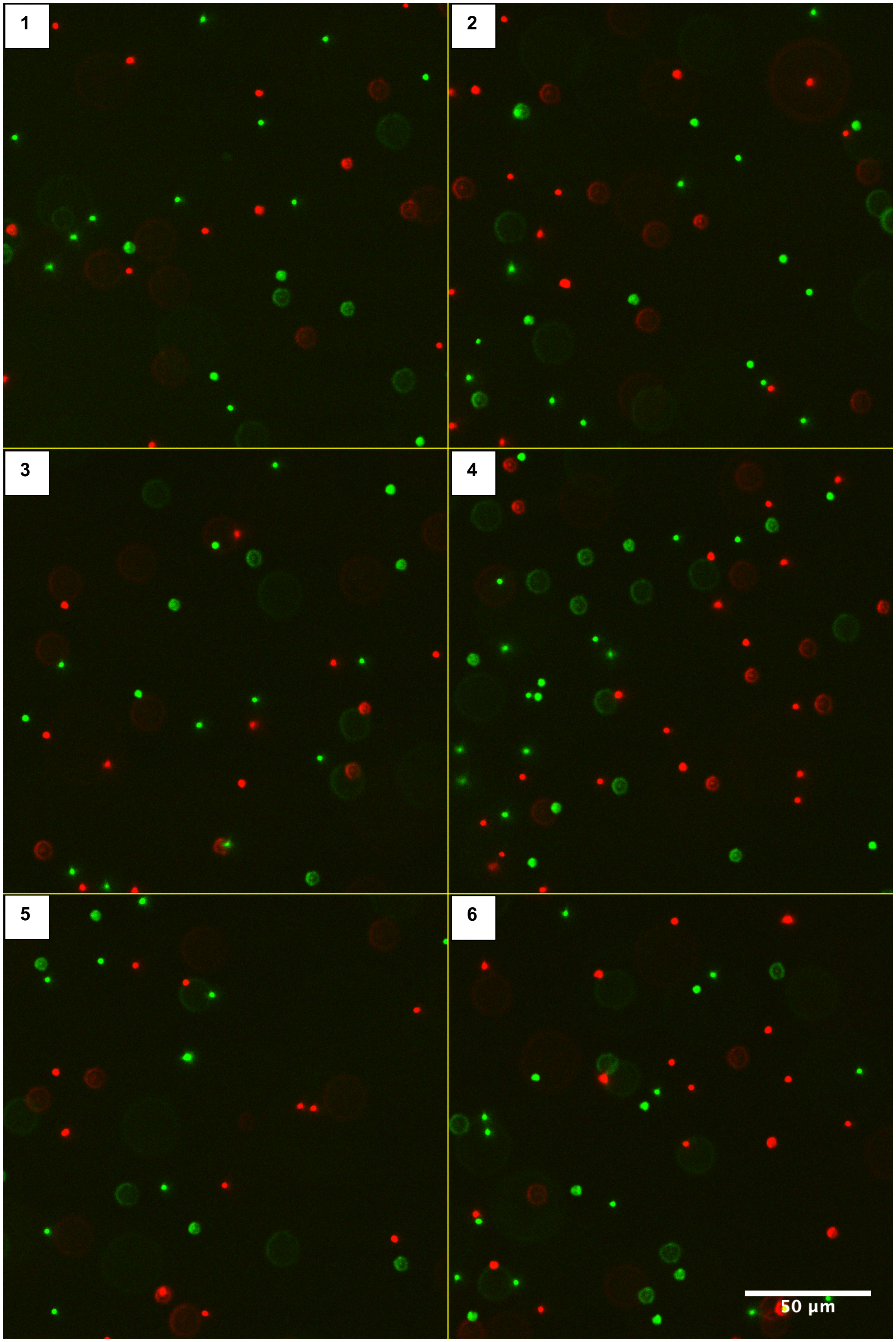

b

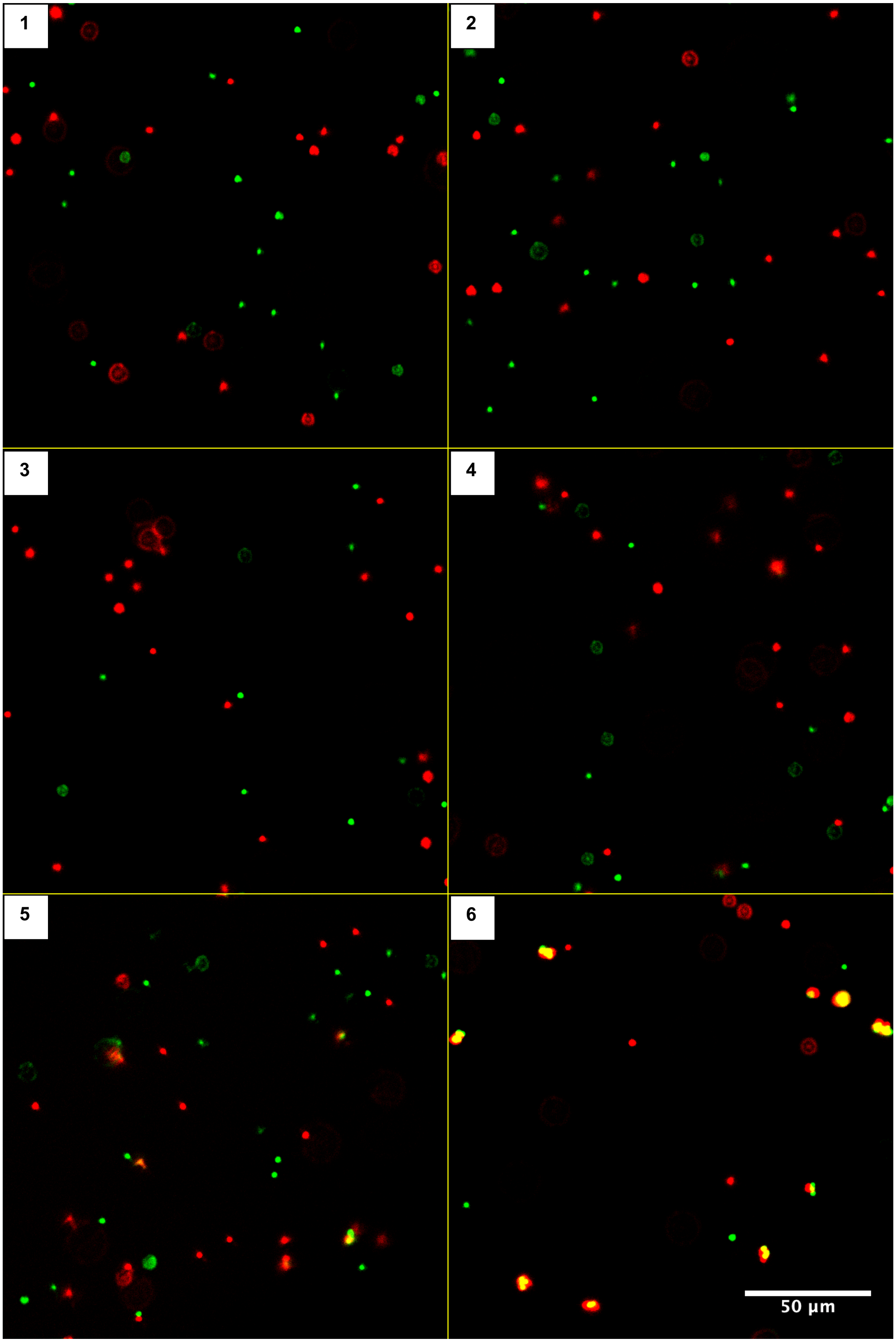

c

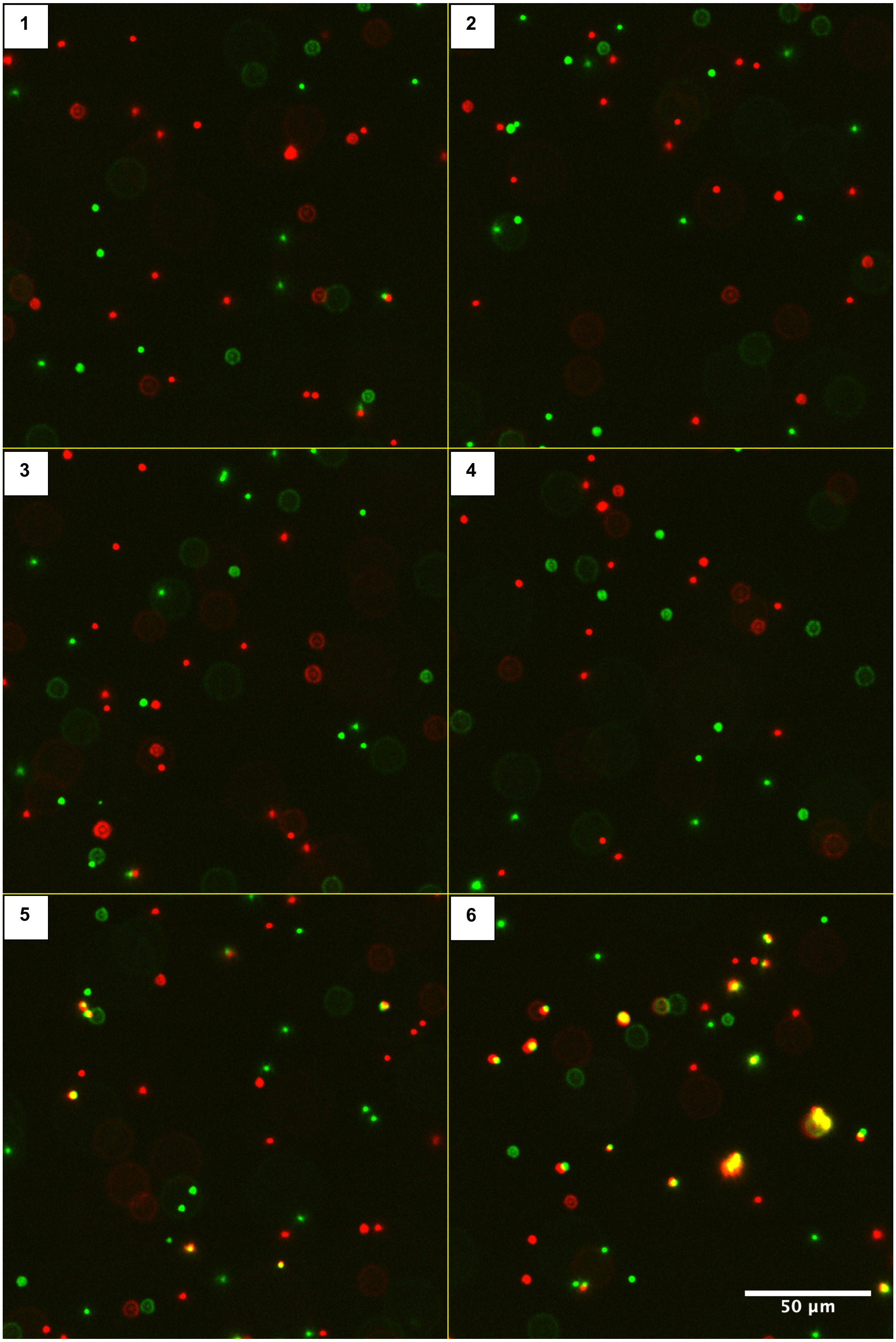

d

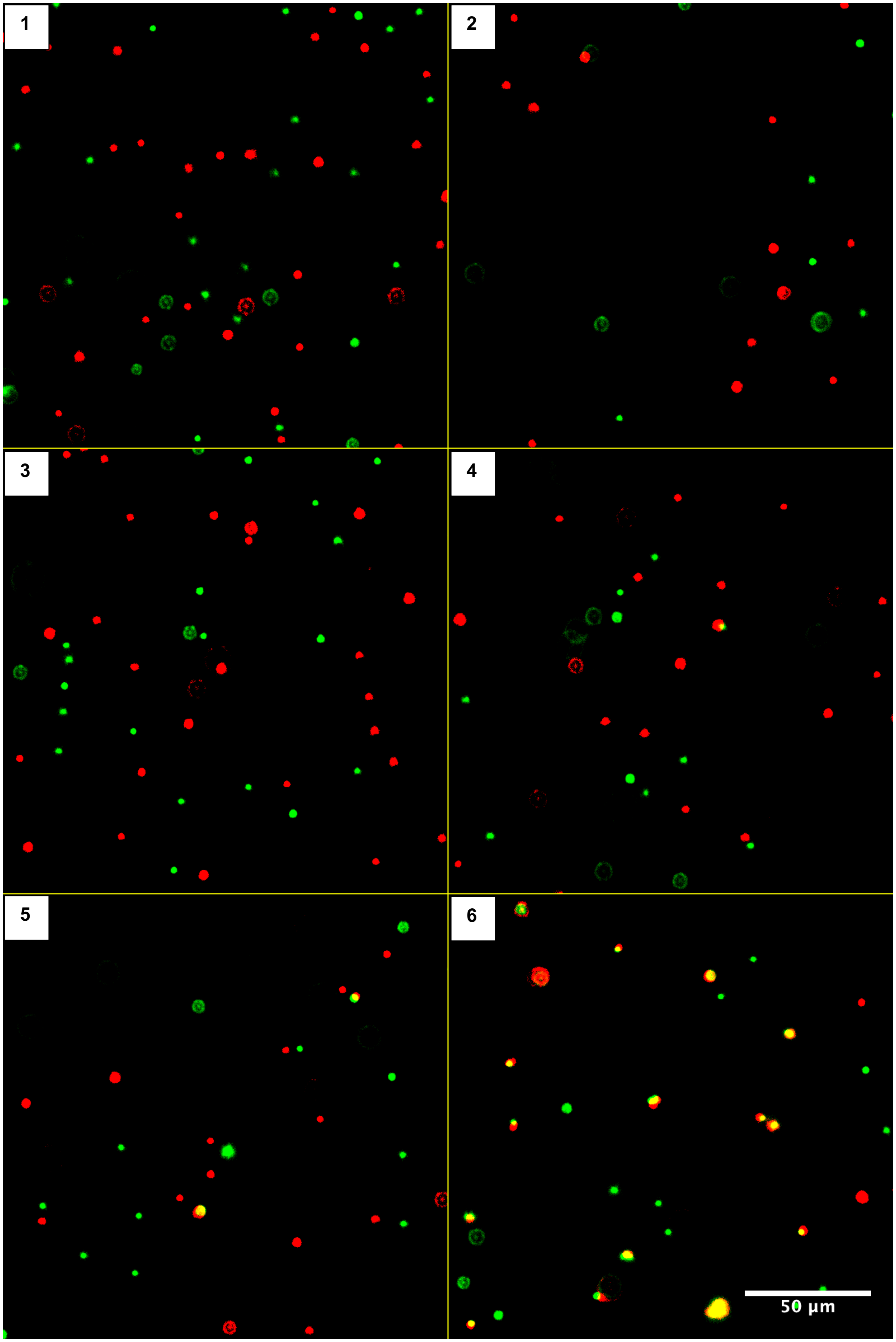

**Figure S2.** Fluorescence images of microbeads at different mixing durations. **(a)** Blank assays comprising six replicates (1–6) mixed for 60 min. **(b–d)** Assays with T1′–2′ mixed for **(b)** 5, **(c)** 30, and **(d)** 60 min. For each mixing duration, six target concentrations were tested: (1) $1.66 \times 10^{-1}$, (2) $1.66 \times 10^{0}$, (3) $1.66 \times 10^{1}$, (4) $1.66 \times 10^{2}$, (5) $1.66 \times 10^{3}$, and (6) $1.66 \times 10^{4}$ fM. Mixing was performed at 400 rpm. G-1 and G-2 microbeads were used at a density of $1.25 \times 10^{3}$ beads/mm$^2$. Assay volume was 50 µL. Images were cropped to 800 × 800 pixels (25% of the original width and height) for presentation. Scale bars, 50 µm.

a

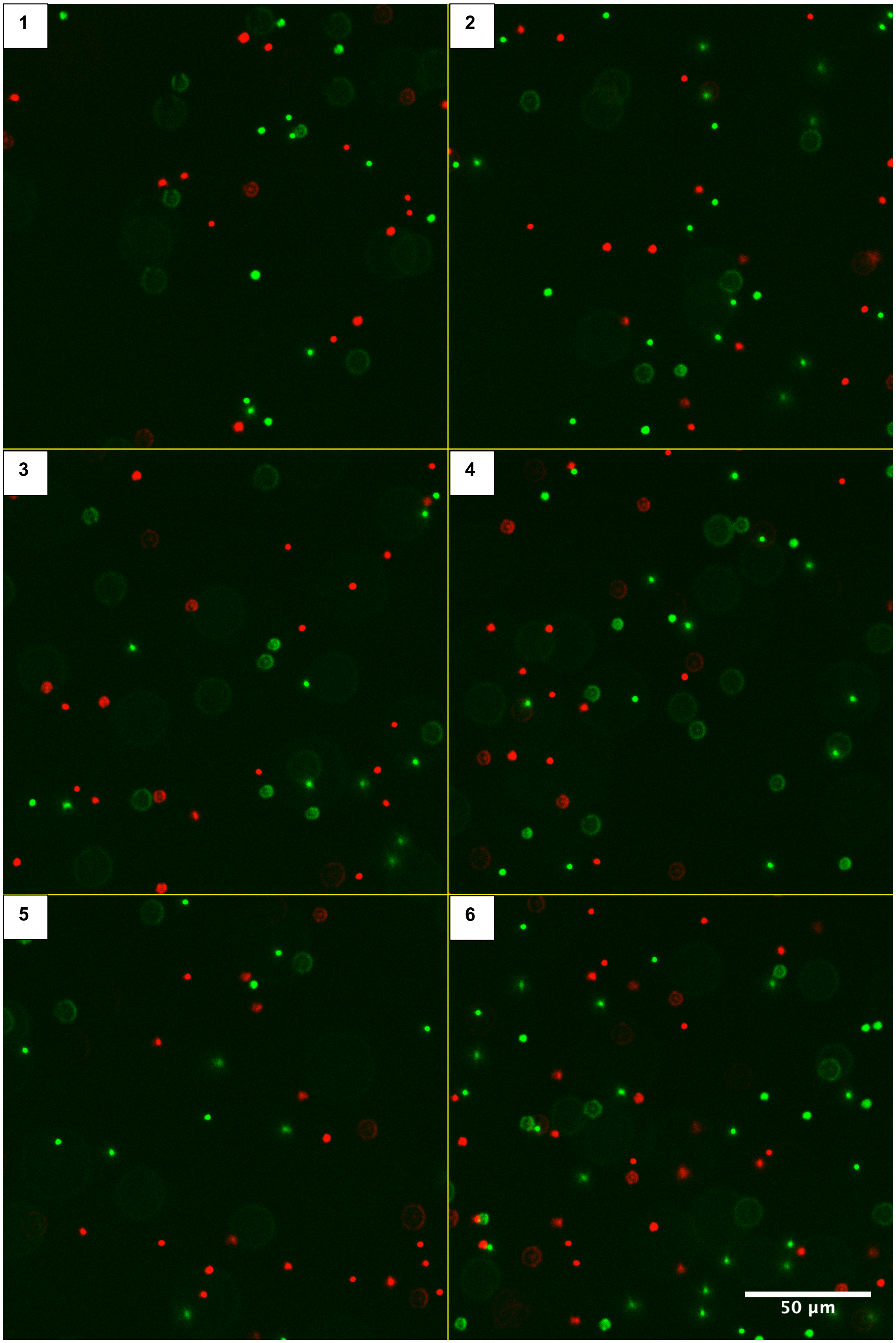

b

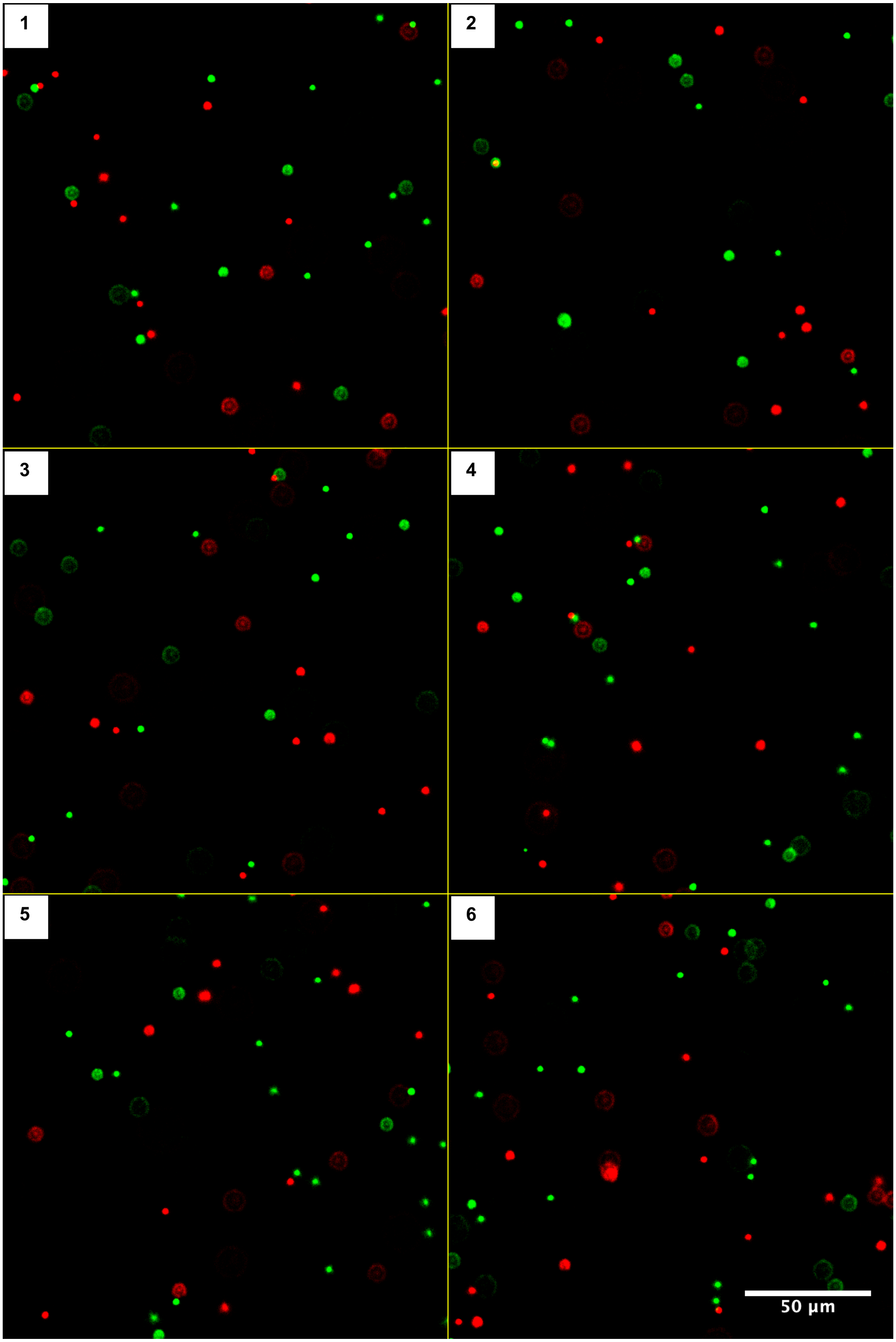

c

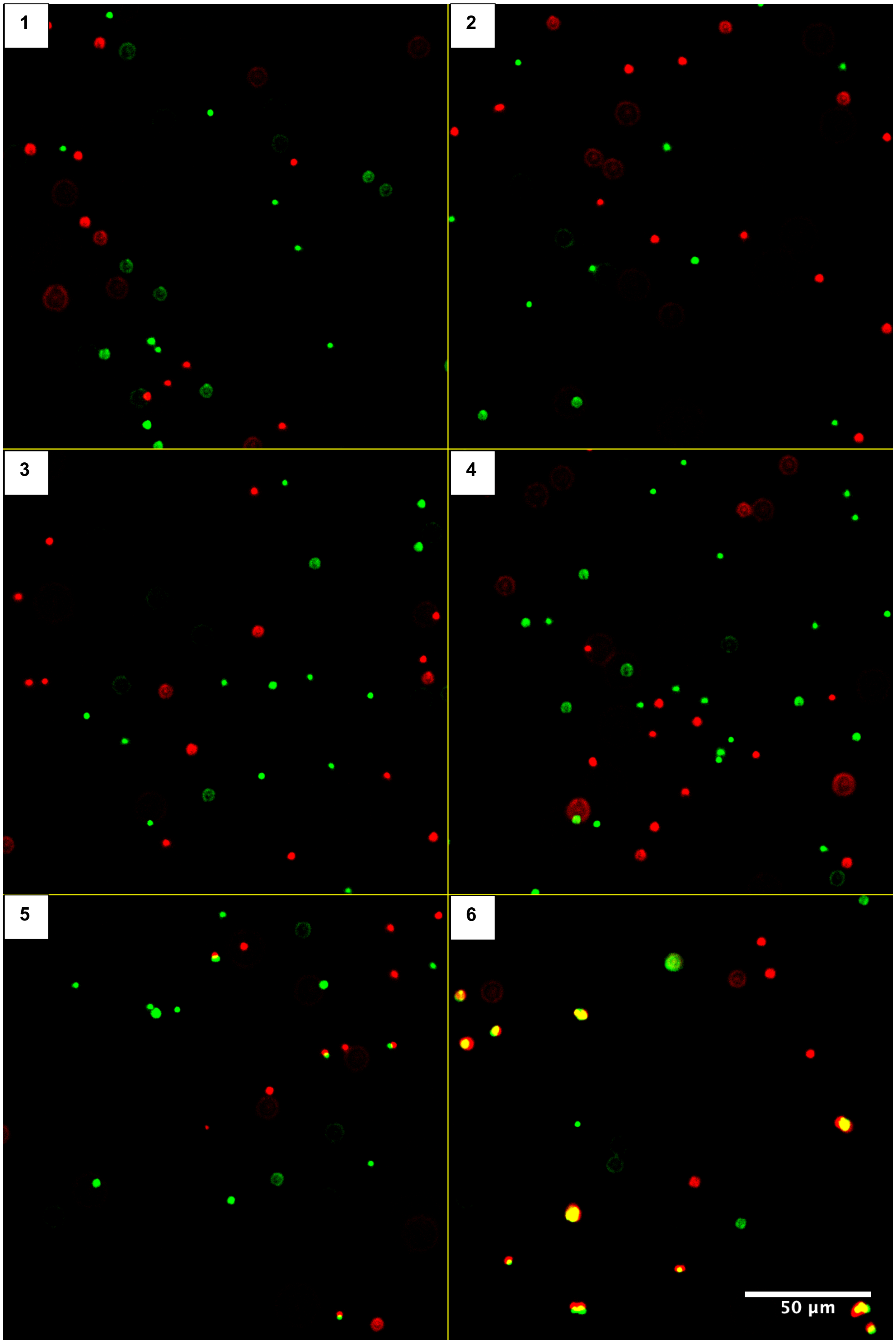

**Figure S3.** Fluorescence images from probe specificity assays. **(a)** Blank assays comprising six replicates (1–6). **(b)** Assays with the scrambled non-target strand Sc1′–2′ at six concentrations: (1) $1.66 \times 10^{-1}$, (2) $1.66 \times 10^{0}$, (3) $1.66 \times 10^{1}$, (4) $1.66 \times 10^{2}$, (5) $1.66 \times 10^{3}$, and (6) $1.66 \times 10^{4}$ fM. **(c)** Assays with target T1′–2′ at the same six concentrations (1–6). G-1 and G-2 microbeads were used at a density of $1.25 \times 10^{3}$ microbeads/mm$^2$ in a 50 μL assay volume. Mixing was performed at 400 rpm for 60 min. Images were cropped to 800 × 800 pixels (25% of the original width and height) for presentation. Scale bars, 50 μm.

a

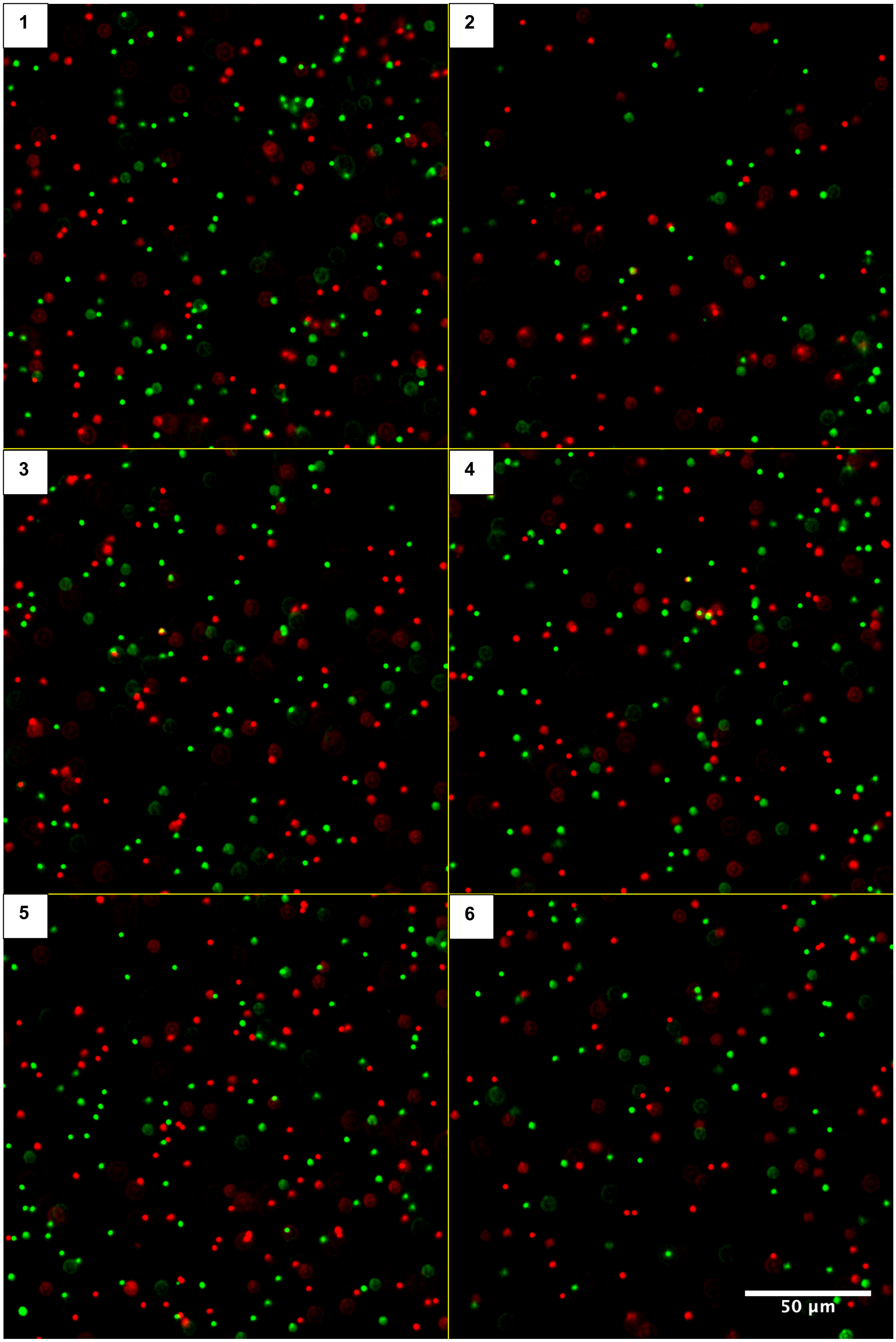

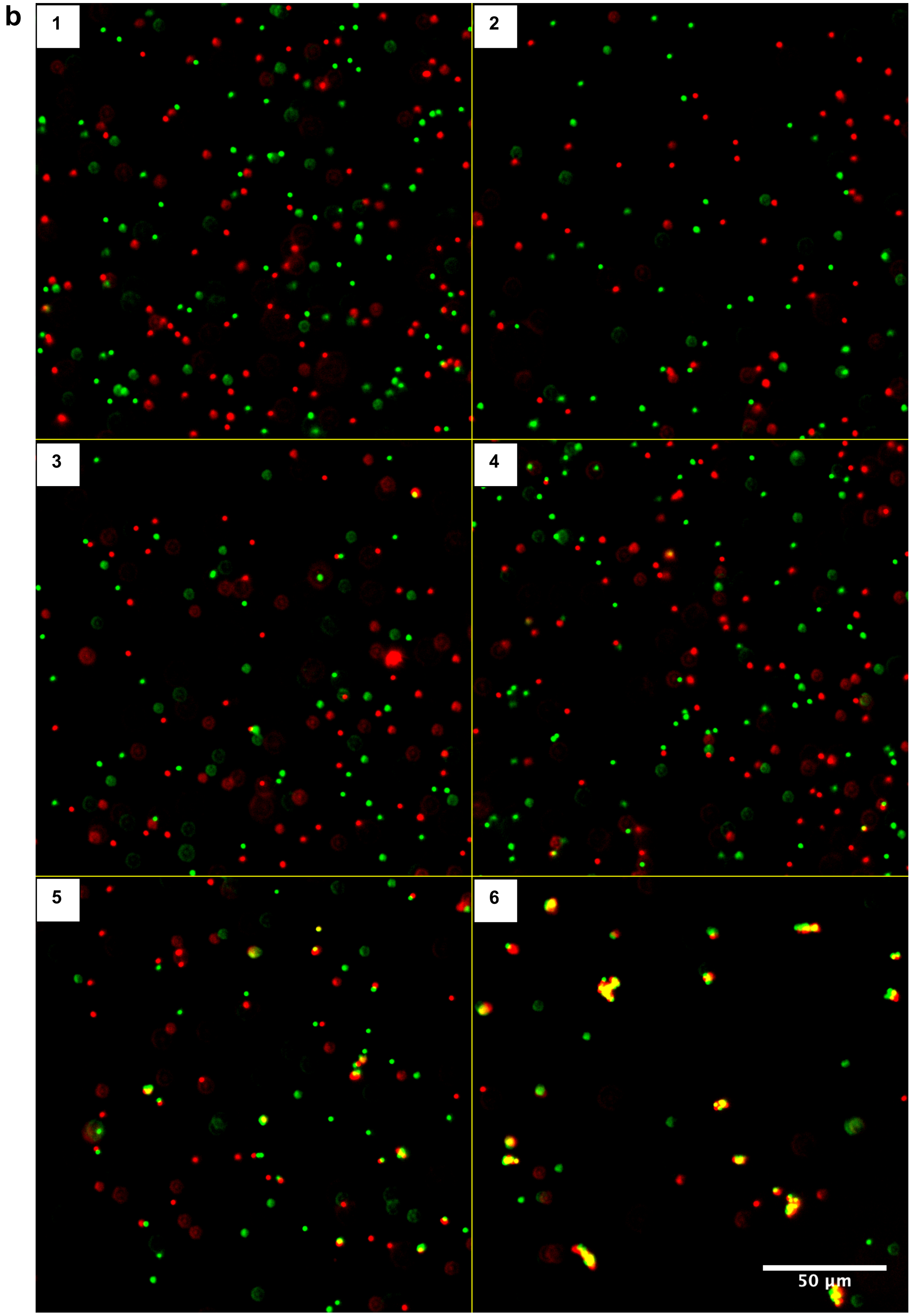
b
1
2
3
4
5
6
50 µm

**Figure S4.** Fluorescence images from the spiking assay. **(a)** Blank assays containing 1 μM Sc1′–2′, comprising six replicates (1–6). **(b)** T1′–2′ with concentrations of (1) $1.66 \times 10^{-1}$, (2) $1.66 \times 10^{0}$, (3) $1.66 \times 10^{1}$, (4) $1.66 \times 10^{2}$, (5) $1.66 \times 10^{3}$, and (6) $1.66 \times 10^{4}$ fM spiked into reaction buffer containing 1 μM Sc1′–2′. G-1 and G-2 microbeads were used at a density of $1.25 \times 10^{3}$ microbeads/mm$^{2}$ in a 25 μL assay volume. Mixing was performed at 400 rpm for 60 min. Images were cropped to 800 × 800 pixels (25% of the original width and height) for presentation. Scale bars, 50 μm.

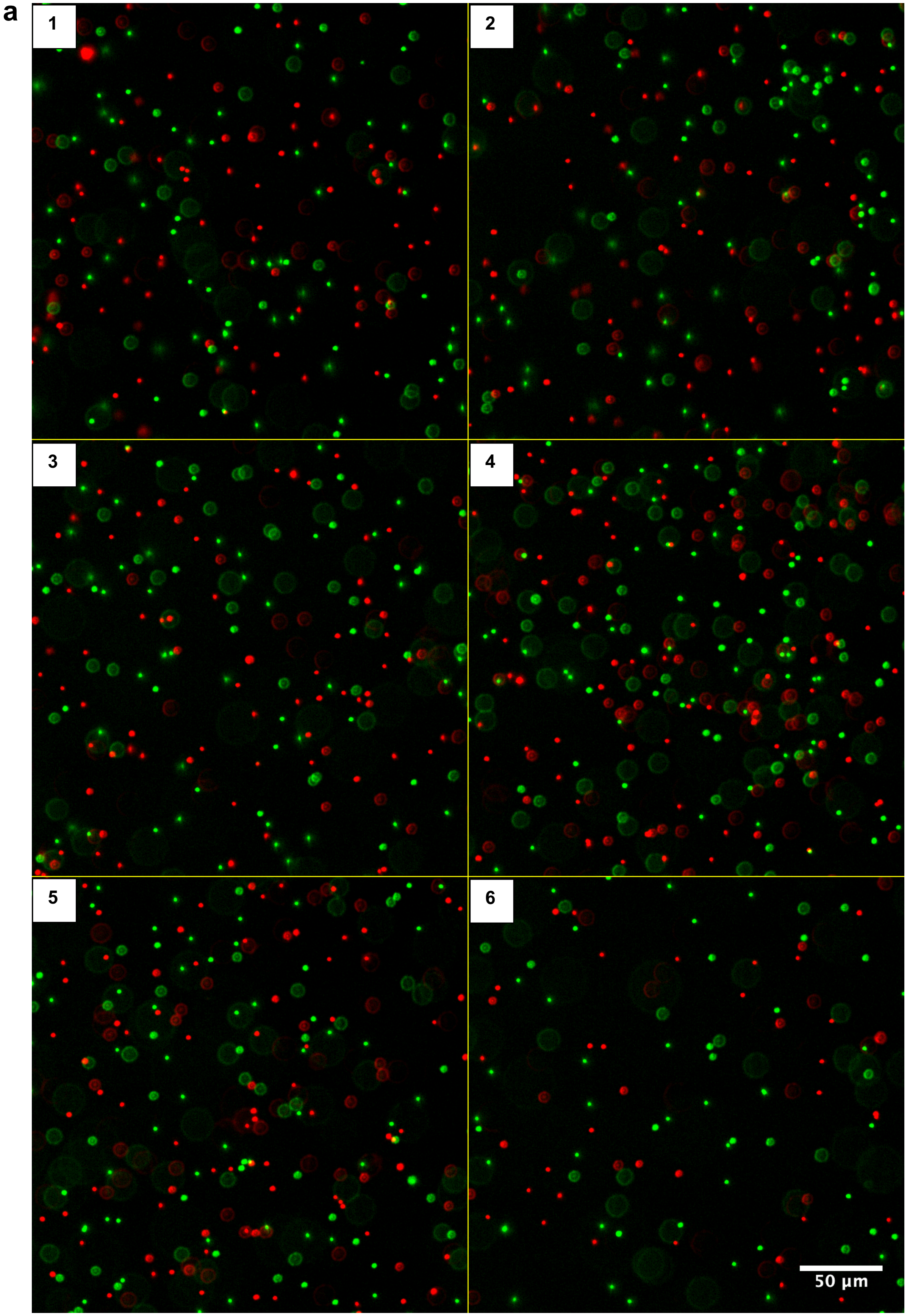
a
1
2
3
4
5
6
50 µm

b

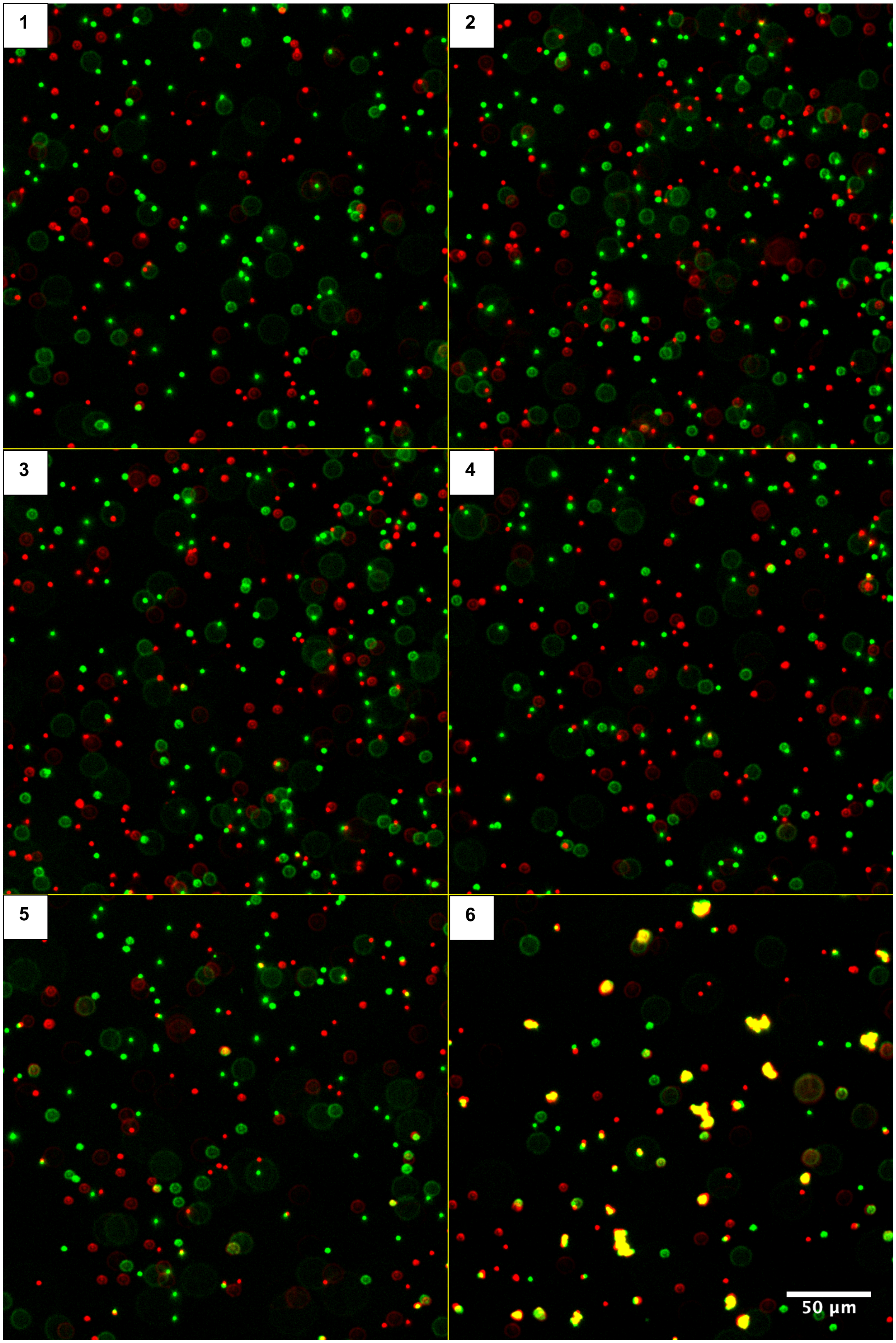

**Figure S5.** Fluorescence images of microbeads without and with SARS-CoV-2-derived synthetic DNA target S3′–4′. **(a)** Blank assays comprising six replicates (1–6). **(b)** Assays with S3′–4′ at six concentrations: (1) $1.66 \times 10^{-1}$, (2) $1.66 \times 10^{0}$, (3) $1.66 \times 10^{1}$, (4) $1.66 \times 10^{2}$, (5) $1.66 \times 10^{3}$, and (6) $1.66 \times 10^{4}$ fM. G-3 and G-4 microbeads were used at a density of $1.25 \times 10^{3}$ microbeads/$mm^2$ in a 50 μL assay volume. Mixing was performed at 400 rpm for 30 min. Images were cropped to 800 × 800 pixels (25% of the original width and height) for presentation. Scale bars, 50 μm.

**Table S1. Oligonucleotides used in this study**

| Name | nt | Sequence (5′–3′) |
|---|---|---|
| P-1 | 35 | Dual biotin/TTCCTCTCACCTCCATCTTACACCACCCACAACAA/Atto488 |
| P-2 | 35 | Atto647N/ACCCTCAAATCAAACCCACCTAACTCCCATCACAC/dual biotin |
| S-1 | 17 | GATGGAGGTGAGAGGAA |
| S-2 | 18 | GTGTGATGGGAGTTAGGT |
| T1′–2′ | 40 | GGTGGGTTTGATTTGAGGGTTTGTTGTGGGTGGTGTAAGA |
| Sc1′–2′ | 70 | CTACGATGTGAGTTAGGCGGGGTTCGCTCGAGCATTTATTTTTGATAGTAC GTGTAGGAGGCGTCAGGAC |
| P-3 | 60 | AAAACGCCTTTTTCAACTTCTACTAAGCCATTGTTGTGGGTGGTGTAAGATG GAGGTGAG |
| P-4 | 60 | ATGGGAGTTAGGTGGGTTTGATTTGAGGGTTCTGCTACCAGCTCAACCATA ACATGACCA |
| S3′–4′ | 60 | TGGCTTAGTAGAAGTTGAAAAAGGCGTTTTTGGTCATGTTATGGTTGAGCT GGTAGCAGA |
| RanN | 25 | NNNNNNNNNNNNNNNNNNNNNNNNN |

nt, nucleotide; P, probe; S, spacer; T1′–2′, target DNA; Sc1′–2′, scrambled non-target DNA; S3′–4′, SARS-CoV-2 ORF1ab-derived synthetic DNA target; RanN, random oligonucleotide, where N denotes A, C, G, or T.

**Table S2.** Heterodimer counts at three different microbead densities, $1.25 \times 10^3$, $2.5 \times 10^3$, and $5 \times 10^3$ microbeads/mm$^2$. Numbers of individual fluorescently labeled microbeads (green for Atto488 and red for Atto647N), heterodimer counts, and heterodimer percentages are shown for blank and T1′–2′-containing assays at each microbead density. The cutoff distance was 6 pixels, corresponding to 1.32 μm using a microscope calibration of 0.22 μm/pixel. The detection threshold was defined as the mean blank signal plus three times the standard deviation of the blank signals. The Z-score was calculated as the difference between the sample signal and the mean blank signal divided by the standard deviation of the blank signals.

| Microbead density | T1′–2′ (fM) | T1′–2′ (copies/μL) | Green bead (G-1) | Red bead (G-2) | Total bead (G-1 & G-2) | Heterodimer count | Heterodimer (%) | Heterodimer *Z*-score | Detection decision (if Z≥3) |
|---|---|---|---|---|---|---|---|---|---|
| | 0 | 0 | 682 | 658 | 1340 | 11 | 1.64 | 1.45 | |
| | 0 | 0 | 653 | 635 | 1288 | 5 | 0.78 | -0.42 | |
| | 0 | 0 | 839 | 862 | 1701 | 12 | 1.41 | 0.95 | |
| | 0 | 0 | 640 | 662 | 1302 | 6 | 0.92 | -0.10 | |
| | 0 | 0 | 649 | 579 | 1228 | 3 | 0.49 | -1.04 | |
| $1.25 \times 10^3$ | 0 | 0 | 532 | 511 | 1043 | 3 | 0.58 | -0.85 | |
| beads/mm$^2$ | **Mean (%)** | | **666** | **651** | **1317** | **7** | **0.97** | | |
| ($5 \times 10^4$ | **Standard deviation, SD (%)** | | | | | | **0.46** | | |
| beads/well) | **Threshold, *T* (%)** | | | | | | **2.36** | | |
| | 1.66E-02 | 1.00E+01 | 505 | 494 | 999 | 7 | 1.40 | 0.93 | Not detected |
| | 1.66E+00 | 1.00E+03 | 453 | 443 | 896 | 2 | 0.45 | -1.12 | Not detected |
| | 1.66E+02 | 1.00E+05 | 460 | 433 | 893 | 18 | 4.03 | 6.61 | Detected |
| | 1.66E+04 | 1.00E+07 | 290 | 283 | 573 | 158 | 55.15 | 116.95 | Detected |
| | **Mean (%)** | | **427** | **413** | **840** | | | | |
| | 0 | 0 | 1431 | 1457 | 2888 | 16 | 1.11 | -1.59 | |
| | 0 | 0 | 1394 | 1379 | 2773 | 37 | 2.67 | 1.47 | |
| | 0 | 0 | 1744 | 1712 | 3456 | 34 | 1.97 | 0.09 | |
| | 0 | 0 | 1702 | 1664 | 3366 | 33 | 1.96 | 0.08 | |
| | 0 | 0 | 1483 | 1536 | 3019 | 26 | 1.72 | -0.39 | |
| $2.5 \times 10^3$ | 0 | 0 | 1569 | 1583 | 3152 | 33 | 2.09 | 0.34 | |
| beads/mm$^2$ | **Mean (%)** | | **1554** | **1555** | **3109** | **30** | **1.92** | | |
| ($1 \times 10^5$ | **Standard deviation, SD (%)** | | | | | | **0.51** | | |
| beads/well) | **Threshold, *T* (%)** | | | | | | **3.45** | | |
| | 1.66E-02 | 1.00E+01 | 1630 | 1648 | 3278 | 39 | 2.38 | 0.90 | Not detected |
| | 1.66E+00 | 1.00E+03 | 1438 | 1497 | 2935 | 29 | 1.98 | 0.11 | Not detected |
| | 1.66E+02 | 1.00E+05 | 1018 | 1057 | 2075 | 39 | 3.76 | 3.61 | Detected |
| | 1.66E+04 | 1.00E+07 | 530 | 565 | 1095 | 346 | 63.20 | 120.15 | Detected |
| | **Mean (%)** | | **1154** | **1192** | **2346** | | | | |
| | 0 | 0 | 1677 | 1715 | 3392 | 48 | 2.83 | 1.45 | |
| | 0 | 0 | 1711 | 1669 | 3380 | 38 | 2.25 | 0.27 | |
| $5 \times 10^3$ | 0 | 0 | 1377 | 1454 | 2831 | 34 | 2.40 | 0.57 | |
| beads/mm$^2$ | 0 | 0 | 1364 | 1363 | 2727 | 27 | 1.98 | -0.29 | |
| ($2 \times 10^5$ | 0 | 0 | 1291 | 1297 | 2588 | 18 | 1.39 | -1.49 | |
| beads/well) | 0 | 0 | 1441 | 1430 | 2871 | 27 | 1.88 | -0.49 | |
| | **Mean (%)** | | **1477** | **1488** | **2965** | **32** | **2.12** | | |
| | **Standard deviation, SD (%)** | | | | | | **0.49** | | |
| | **Threshold, *T* (%)** | | | | | | **3.60** | | |
| | 1.66E-02 | 1.00E+01 | 1599 | 1616 | 3215 | 30 | 1.87 | -0.51 | Not detected |
| | 1.66E+00 | 1.00E+03 | 1507 | 1403 | 2910 | 35 | 2.41 | 0.59 | Not detected |
| | 1.66E+02 | 1.00E+05 | 1554 | 1503 | 3057 | 98 | 6.41 | 8.76 | Detected |
| | 1.66E+04 | 1.00E+07 | 935 | 925 | 1860 | 638 | 68.60 | 135.67 | Detected |
| | **Mean (%)** | | **1399** | **1362** | **2761** | | | | |

**Table S3.** Heterodimer counts after 5, 30, and 60 min of mixing. Numbers of individual fluorescently labeled microbeads, heterodimer counts, and heterodimer percentages are shown for T1′–2′-containing assays at each mixing duration. Blank assays without T1′–2′ were performed only under the 60 min mixing condition. The cutoff distance was 8 pixels, corresponding to 1.76 µm using a microscope calibration of 0.22 µm/pixel. The detection threshold and Z-score were calculated as described for Table S2.

| Mixing duration | T1′–2′ (fM) | T1′–2′ (copies/µL) | Green bead (G-1) | Red bead (G-2) | Total bead (G-1 & G-2) | Heterodimer count | Heterodimer (%) | Heterodimer $Z$-score | Detection decision (if Z≥3) |
|---|---|---|---|---|---|---|---|---|---|
| 60 min | 0 | 0 | 469 | 358 | 827 | 5 | 1.21 | 0.50 | |
| | 0 | 0 | 480 | 428 | 908 | 3 | 0.66 | -0.67 | |
| | 0 | 0 | 368 | 333 | 701 | 2 | 0.57 | -0.87 | |
| | 0 | 0 | 382 | 369 | 751 | 2 | 0.53 | -0.95 | |
| | 0 | 0 | 376 | 325 | 701 | 6 | 1.71 | 1.56 | |
| | 0 | 0 | 334 | 342 | 676 | 4 | 1.18 | 0.43 | |
| | **Mean (%)** | | **402** | **359** | **761** | **4** | **0.98** | | |
| | **Standard deviation, SD (%)** | | | | | | **0.47** | | |
| | **Threshold, *T* (%)** | | | | | | **2.38** | | |
| 5 min | 1.66E-01 | 1.00E+02 | 411 | 359 | 770 | 7 | 1.82 | 1.79 | Not detected |
| | 1.66E+00 | 1.00E+03 | 432 | 378 | 810 | 2 | 0.49 | -1.04 | Not detected |
| | 1.66E+01 | 1.00E+04 | 400 | 376 | 776 | 5 | 1.29 | 0.67 | Not detected |
| | 1.66E+02 | 1.00E+05 | 403 | 358 | 761 | 6 | 1.58 | 1.28 | Not detected |
| | 1.66E+03 | 1.00E+06 | 429 | 390 | 819 | 59 | 14.41 | 28.58 | Detected |
| | 1.66E+04 | 1.00E+07 | 176 | 176 | 352 | 109 | 61.93 | 129.69 | Detected |
| | **Mean (%)** | | **375** | **340** | **715** | **31** | | | |
| 30 min | 1.66E-01 | 1.00E+02 | 411 | 382 | 793 | 7 | 1.77 | 1.69 | Not detected |
| | 1.66E+00 | 1.00E+03 | 407 | 380 | 787 | 3 | 0.76 | -0.46 | Not detected |
| | 1.66E+01 | 1.00E+04 | 392 | 365 | 757 | 4 | 1.06 | 0.18 | Not detected |
| | 1.66E+02 | 1.00E+05 | 360 | 326 | 686 | 12 | 3.50 | 5.37 | Detected |
| | 1.66E+03 | 1.00E+06 | 450 | 366 | 816 | 53 | 12.99 | 25.56 | Detected |
| | 1.66E+04 | 1.00E+07 | 382 | 384 | 766 | 239 | 62.40 | 130.69 | Detected |
| | **Mean (%)** | | **400** | **367** | **768** | **53** | | | |
| 60 min | 1.66E-01 | 1.00E+02 | 482 | 485 | 967 | 4 | 0.83 | -0.31 | Not detected |
| | 1.66E+00 | 1.00E+03 | 352 | 329 | 681 | 3 | 0.88 | -0.21 | Not detected |
| | 1.66E+01 | 1.00E+04 | 338 | 352 | 690 | 6 | 1.74 | 1.62 | Not detected |
| | 1.66E+02 | 1.00E+05 | 379 | 374 | 753 | 14 | 3.72 | 5.84 | Detected |
| | 1.66E+03 | 1.00E+06 | 365 | 339 | 704 | 49 | 13.92 | 27.54 | Detected |
| | 1.66E+04 | 1.00E+07 | 286 | 271 | 557 | 160 | 57.45 | 120.16 | Detected |
| | **Mean (%)** | | **367** | **358** | **725** | **39** | | | |

**Table S4.** Heterodimer counts from probe specificity assays with no added DNA (blank), the scrambled non-target strand Sc1′–2′, or target T1′–2′. The cutoff distance was 6 pixels, corresponding to 1.32 µm using a microscope calibration of 0.22 µm/pixel. The detection threshold and Z-score were calculated as described for Table S2.

| Assay | Target (fM) | Target (copies/µL) | Green bead (G-1) | Red bead (G-2) | Total bead (G-1 & G-2) | Heterodimer count | Heterodimer (%) | Heterodimer *Z*-score | Detection decision (if Z≥3) |
|---|---|---|---|---|---|---|---|---|---|
| | 0 | 0 | 316 | 327 | 643 | 0 | 0.00 | -1.21 | |
| | 0 | 0 | 345 | 292 | 637 | 2 | 0.63 | 0.44 | |
| | 0 | 0 | 387 | 320 | 707 | 0 | 0.00 | -1.21 | |
| | 0 | 0 | 448 | 404 | 852 | 2 | 0.47 | 0.03 | |
| Blank | 0 | 0 | 375 | 337 | 712 | 3 | 0.84 | 1.01 | |
| | 0 | 0 | 530 | 452 | 982 | 4 | 0.81 | 0.93 | |
| | **Mean (%)** | | **400** | **355** | **756** | **2** | **0.46** | | |
| | **Standard deviation, SD (%)** | | | | | | **0.38** | | |
| | **Threshold, *T* (%)** | | | | | | **1.60** | | |
| | 1.66E-01 | 1.00E+02 | 348 | 326 | 674 | 1 | 0.30 | -0.42 | Not detected |
| | 1.66E+00 | 1.00E+03 | 351 | 302 | 653 | 2 | 0.61 | 0.40 | Not detected |
| | 1.66E+01 | 1.00E+04 | 368 | 294 | 662 | 3 | 0.91 | 1.19 | Not detected |
| Sc1′–2′ | 1.66E+02 | 1.00E+05 | 358 | 305 | 663 | 4 | 1.21 | 1.97 | Not detected |
| | 1.66E+03 | 1.00E+06 | 369 | 313 | 682 | 3 | 0.88 | 1.11 | Not detected |
| | 1.66E+04 | 1.00E+07 | 421 | 386 | 807 | 4 | 0.99 | 1.40 | Not detected |
| | **Mean (%)** | | **369** | **321** | **690** | | **0.82** | | |
| | 1.66E-01 | 1.00E+02 | 387 | 336 | 723 | 3 | 0.83 | 0.97 | Not detected |
| | 1.66E+00 | 1.00E+03 | 396 | 310 | 706 | 4 | 1.13 | 1.76 | Not detected |
| | 1.66E+01 | 1.00E+04 | 388 | 331 | 719 | 1 | 0.28 | -0.47 | Not detected |
| T1′–2′ | 1.66E+02 | 1.00E+05 | 385 | 353 | 738 | 16 | 4.34 | 10.20 | Detected |
| | 1.66E+03 | 1.00E+06 | 346 | 305 | 651 | 61 | 18.74 | 48.05 | Detected |
| | 1.66E+04 | 1.00E+07 | 235 | 207 | 442 | 89 | 40.27 | 104.64 | Detected |
| | **Mean (%)** | | **356** | **307** | **663** | | | | |

**Table S5.** Heterodimer counts from spiking assays. Blank assays contained 1 μM scrambled non-target strand Sc1′–2′, and six concentrations of T1′–2′ were spiked into the same non-target background. The cutoff distance was 6 pixels, corresponding to 1.32 μm using a microscope calibration of 0.22 μm/pixel. The detection threshold and Z-score were calculated as described for Table S2.

| Assay | T1′–2′ (fM) | T1′–2′ (copies/μL) | Green bead (G-1) | Red bead (G-2) | Total bead (G-1 & G-2) | Heterodimer count | Heterodimer (%) | Heterodimer *Z*-score | Detection decision (if Z≥3) |
|---|---|---|---|---|---|---|---|---|---|
| Blank with 1 μM of Sc1′–2′ | 0 | 0 | 1316 | 1368 | 2684 | 25 | 1.86 | 0.20 | |
| | 0 | 0 | 911 | 917 | 1828 | 21 | 2.30 | 1.30 | |
| | 0 | 0 | 1274 | 1230 | 2504 | 22 | 1.76 | -0.07 | |
| | 0 | 0 | 1273 | 1267 | 2540 | 25 | 1.97 | 0.47 | |
| | 0 | 0 | 1077 | 1127 | 2204 | 19 | 1.72 | -0.16 | |
| | 0 | 0 | 1073 | 1106 | 2179 | 12 | 1.10 | -1.74 | |
| | **Mean (%)** | | **1154** | **1169** | **2323** | **21** | **1.79** | | |
| | **Standard deviation, SD (%)** | | | | | | **0.39** | | |
| | **Threshold, *T* (%)** | | | | | | **2.97** | | |
| T1′–2′ spiked in 1 μM of Sc1′–2′ | 1.66E-01 | 1.00E+02 | 1233 | 1321 | 2554 | 22 | 1.72 | -0.16 | Not detected |
| | 1.66E+00 | 1.00E+03 | 1019 | 1077 | 2096 | 7 | 0.67 | -2.84 | Not detected |
| | 1.66E+01 | 1.00E+04 | 908 | 997 | 1905 | 16 | 1.68 | -0.27 | Not detected |
| | 1.66E+02 | 1.00E+05 | 1175 | 1237 | 2412 | 57 | 4.73 | 7.47 | Detected |
| | 1.66E+03 | 1.00E+06 | 750 | 682 | 1432 | 106 | 14.80 | 33.08 | Detected |
| | 1.66E+04 | 1.00E+07 | 350 | 352 | 702 | 180 | 51.28 | 125.78 | Detected |
| | **Mean (%)** | | **906** | **944** | **1850** | | | | |

**Table S6.** Heterodimer counts for the SARS-CoV-2 ORF1ab-derived synthetic DNA target S3′–4′. Assays contained G-3/G-4 microbeads at $1.25 \times 10^3$ microbeads/mm$^2$. The cutoff distance was 6 pixels, corresponding to 1.98 µm using a microscope calibration of 0.33 µm/pixel. The detection threshold and Z-score were calculated as described for Table S2.

| Assay | S3′–4′ (fM) | S3′–4′ (copies/µL) | Green bead (G-3) | Red bead (G-4) | Total bead (G-3 & G-4) | Heterodimer count | Heterodimer (%) | Heterodimer *Z*-score | Detection decision (if Z≥3) |
|---|---|---|---|---|---|---|---|---|---|
| Blank | 0 | 0 | 1231 | 1343 | 2574 | 16 | 1.24 | -0.99 | |
| | 0 | 0 | 1166 | 1260 | 2426 | 30 | 2.47 | 1.38 | |
| | 0 | 0 | 1218 | 1311 | 2529 | 20 | 1.58 | -0.34 | |
| | 0 | 0 | 1375 | 1503 | 2878 | 30 | 2.08 | 0.63 | |
| | 0 | 0 | 1250 | 1342 | 2592 | 26 | 2.01 | 0.48 | |
| | 0 | 0 | 933 | 982 | 1915 | 11 | 1.15 | -1.17 | |
| | **Mean (%)** | | **1196** | **1290** | **2486** | **22** | **1.76** | | |
| | **Standard deviation, SD (%)** | | | | | | **0.52** | | |
| | **Threshold, *T* (%)** | | | | | | **3.31** | | |
| S3′–4′ | 1.66E-01 | 1.00E+02 | 1481 | 1615 | 3096 | 33 | 2.13 | 0.72 | Not detected |
| | 1.66E+00 | 1.00E+03 | 1653 | 1733 | 3386 | 38 | 2.24 | 0.93 | Not detected |
| | 1.66E+01 | 1.00E+04 | 1845 | 2094 | 3939 | 65 | 3.3 | 2.97 | Not detected |
| | 1.66E+02 | 1.00E+05 | 1449 | 1540 | 2989 | 66 | 4.42 | 5.13 | Detected |
| | 1.66E+03 | 1.00E+06 | 1294 | 1360 | 2654 | 323 | 24.34 | 43.51 | Detected |
| | 1.66E+04 | 1.00E+07 | 617 | 649 | 1266 | 333 | 52.61 | 97.97 | Detected |
| | **Mean (%)** | | **1390** | **1499** | **2888** | | | | |

**Mathematica Code S1.** Image-based microbead and heterodimer quantification script

```
(*1. get the file names*)
SetDirectory@NotebookDirectory[];
files = Sort@FileNames["*STACK.tif", NotebookDirectory[], 3];
(*2. set the quantile for adjustment of image intensity*)
quant = {#, 1 - #} &@0.001;
(*3. import all images and arrange them as two-channel image pairs*)
sx = ArrayReshape[#, {Length[#]/2, 2}] &[
  ImageAdjust[#, 0, Quantile[Flatten@ImageData@#, quant]] & /@ Flatten[Import /@ files]];
(*For these datasets: channel 1 = Atto488 (green); channel 2 = Atto647N (red).*)
greenChannel = 1; redChannel = 2;
(*4. determine centroid coordinates of detected fluorescent components*)
coord = ParallelMap[
  ComponentMeasurements[#, "Centroid"][[All, 2]] &@
    MorphologicalComponents@MaxDetect[Erosion[#, 1], 0.1] & /@ # &, sx];
(*5. visually check component detection*)
nf = Map[Nearest, coord, {2}];
index = 5; (*change to inspect a different image*)
radius = 7;
imagetest = Show[
  ColorCombine[{#[[redChannel]], #[[greenChannel]], Image@ConstantArray[0,
ImageDimensions@#[[1]]]}, "RGB"] &@ sx[[index]],
  Graphics[{Red, Circle[#, radius]} & /@ coord[[index, redChannel]]],
  Graphics[{Green, Circle[#, radius]} & /@ coord[[index, greenChannel]]]];
distance = 6; (*experiment-specific cutoffs ranged from 6 to 8 pixels*)
controlwell = 6; (*number of blank wells*)
factor = 1;
targetConcentrations = {0.166, 1.66, 16.6, 166, 1660, 16600}; (*edit for each experiment*)
concentration = Join[ConstantArray[0, controlwell], targetConcentrations];
assayName = "Assay name";
(*6. heterodimer quantification*)
dat = ParallelTable[Module[{green, red, distanceLists, heterodimers},
  green = factor*Length@coord[[index, greenChannel]];
  red = factor*Length@coord[[index, redChannel]];
```

```
  distanceLists = Map[Norm, nf[[index, greenChannel]][#, 30] - ConstantArray[#, 30]] & /@
coord[[index, redChannel]];
  heterodimers = factor*Count[Flatten@distanceLists, n_ /; n <= distance];
  {index, concentration[[index]], green, red, heterodimers, 100*N@heterodimers/Mean[{green, red}]}],
  {index, Length@sx}];
{mean, std} = {Mean@#, StandardDeviation@#} &@dat[[;; controlwell, -1]];
threshold = mean + 3*std;
datNormalize = Append[#, (#[[-1]] - mean)/std] & /@ dat;
(*7. display results in a table*)
TableForm[datNormalize, TableHeadings -> {None,
  {"sample", "fM", "# of green beads", "# of red beads", "# of heterodimers", "Heterodimer %", "z
score"}}];
(*8. display the target-response graph*)
targetWells = Range[controlwell + 1, Length@concentration];
Show[ListLogLogPlot[Table[{concentration[[well]], dat[[well, -1]]}, {well, targetWells}],
  Frame -> True, FrameLabel -> {"Target concentration (fM)", "Heterodimer percentage"}, PlotLabel ->
assayName],
 LogLogPlot[threshold, {x, 0.00001, 10^8}, PlotStyle -> {Red}],
 ImageSize -> Large, LabelStyle -> {12, GrayLevel[0]}];
(*9. export results as CSV*)
headers = {"sample", "fM", "# of green beads", "# of red beads", "# of heterodimers", "Heterodimer %",
"z score"};
outputFile = SystemDialogInput["FileSave", "heterodimer_quantification.csv"];
If[outputFile =!= $Canceled, Export[outputFile, Prepend[datNormalize, headers], "CSV"];
  Print["Saved to: ", outputFile];]
```
